# Intrinsic cardiac adrenergic cells contribute to septic cardiomyopathy

**DOI:** 10.1101/2021.03.02.433552

**Authors:** Duomeng Yang, Xiaomeng Dai, Yun Xing, Xiangxu Tang, Guang Yang, Penghua Wang, Andrew G. Harrison, Hongmei Li, Xiuxiu Lv, Xiaohui Yu, Huadong Wang

## Abstract

Occurring independently of cardiac sympathetic nervous system, the intrinsic cardiac adrenergic (ICA) cells have been identified as an important regulator in both of developing and adult cardiac physiological and pathological processes. However, its role in septic cardiomyopathy remains unknown. Herein, we report that lipopolysaccharide (LPS) dose- and time-dependently increased norepinephrine (NE) release from ICA cells, which aggravates myocardial TNF-α production and dysfunction. Inhibition of NE synthesis in ICA cells alleviated LPS-elicited cardiac dysfunction as well as TNF-α production in Langendorff perfusing hearts. Mechanistically, ICA cell expressed Toll-like receptor 4 (TLR4), activated by LPS, to increase the expression of tyrosine hydroxylase, a key enzyme responsible for NE biosynthesis, via AP-1 binding to its promoter. Surprisingly, LPS-TLR4 signaling triggered no TNF-α production in ICA cells due to the elevated *Nfkbia* and *Tnfaip6* expression. In LPS-treated co-culture of ICA cells and cardiomyocytes, the raised NE from ICA cells activated cardiomyocyte β_1_-adrenergic receptor (β_1_-AR), driving Ca^2+^/calmodulin-dependent protein kinase II (CaMKII) to increase the activities of NF-κB and mitogen-activated protein kinase pathways, which were mimicked by dobutamine. Our findings reveal a cell type-specific TLR4 function triggering NE synthesis, but not TNF-α production in inflammatory pathogenesis, and identify ICA cell-derived NE as a paracrine signal in the cross talk among different cardiac cells to enhance myocardial injury during LPS challenge, suggesting that targeting ICA cell-derived NE may be a potential therapeutic strategy for septic cardiomyopathy.

## Introduction

Myocardial dysfunction is a frequent event that corresponds to the severity of sepsis, which accounts for the primary cause of death in intensive care units.^1,2^ During sepsis-induced myocardial dysfunction (SIMD), pathogen-associated molecular patterns, such as bacterial lipopolysaccharide (LPS), interact with Toll-like receptor 4 (TLR4) on immune and cardiac cells, activating NF-κB and mitogen-activated protein kinase (MAPK) signaling cascades and causing a burst of proinflammatory cytokines, including TNF-α, intrleukin-1β and intrleukin-6. These cytokines cause cardiac depression directly.^3-8^ Norepinephrine (NE), demonstrated to be associated with SIMD,^9,10^ can activate β_1_-adrenergic receptor (AR) and promote Ca^2+^/calmodulin-dependent protein kinase II (CaMKII) phosphorylation, IκBα phosphorylation as well as TNF-α expression, subsequently contribute to cardiomyocyte apoptosis in septic mice.^11^ Moreover, selective β_1_-AR blockade improves cardiac function and survival in septic patients.^12,13^

The intrinsic cardiac adrenergic (ICA) cells, which express tyrosine hydroxylase (TH) and dopamine-β-hydroxylase (DBH) necessary for catecholamine biosynthesis, have been identified in the mammalian heart as an important origin of intrinsic cardiac catecholamines.^14,15^ A growing body of literature has proven that the ICA system is a critical regulator of mammalian heart development, post-heart transplantation inotropic support and cardioprotection during ischemia/reperfusion.^15-21^ However, the response of ICA cells in sepsis is unclear and the role of ICA cells in SIMD is not established. Our previous study showed that blockade of β_1_-AR suppressed LPS-induced TNF-α production in primary neonatal rat cardiomyocytes cultured by traditional methods in the absence of exogenous NE,^22^ indicating that some β_1_-AR stimuli exist in the primary neonatal rat cardiomyocyte culture obtained by traditional methods. In addition, ICA cells were found to be present in the primary neonatal rat cardiomyocyte culture obtained by traditional methods.^14,23^ In this context, we postulated that LPS might stimulate ICA cells to synthesize NE and the ICA cell-derived NE would promote LPS-induced cardiomyocyte TNF-α production and myocardial dysfunction via β_1_-AR.

In this study, we investigated the effect of LPS on NE production in ICA cells and underlying mechanisms, and determined the role of ICA cell-derived NE in LPS-induced cardiomyocyte TNF-α production and myocardial dysfunction. We first demonstrated, as far as we know, that LPS increased NE production in ICA cells that express TLR4, and identified ICA cell-derived NE as a paracrine signal to enhance LPS-provoked proinflammatory response in cardiomyocytes and aggravate LPS-induced myocardial dysfunction. These findings reveal a pivotal role of ICA cells in the development of sepsis-induced myocardial dysfunction.

## Results

### ICA cells, as mesenchymal cells, exist in the primary neonatal rat ventricular myocyte culture obtained by traditional enzymatic methods and adult rat hearts

As a key rate-limiting enzyme for catecholamine biosynthesis, TH was used as a marker of ICA cells (Fig. S1B).^17^ As shown in Fig. 1a, TH-positive ICA cells were found to be growing in clusters along with TH-negative neonatal rat ventricular myocytes (NRVM) which are cardiac troponin I (cTn I)-positive isolated by traditional enzymatic methods described previously,^24^ and these ICA cells expressing TH were found in adult rat hearts as well (Fig. 1b). In addition, the established 3D structure of ICA cells further illustrated its TH expression (Fig. S1C). Macrophages have been reported as a source of catecholamines in response to acute inflammatory injury,^25^ but we demonstrated that ICA cells did not express macrophage markers CD68 or F4/80 (Fig. 1c and Fig. S1D, E). Moreover, single-cell RNA sequencing revealed ICA cells and macrophages as two distinct clusters (Fig. 5d, e) with upregulation of *Dbh* expression in ICA cells and a deficiency in macrophages following LPS stimulation (Fig. 5h, i). Therefore, ICA cells were not considered to be of the macrophage lineage. Since vimentins are type III intermediate filaments expressed especially in mesenchymal cells, co-stainning of cTn I, TH and vimentin in co-culture of ICA cells and myocytes demonstrated that ICA cells were mesenchymal cells in rat hearts (Fig. 1d).

**Fig. 1.**
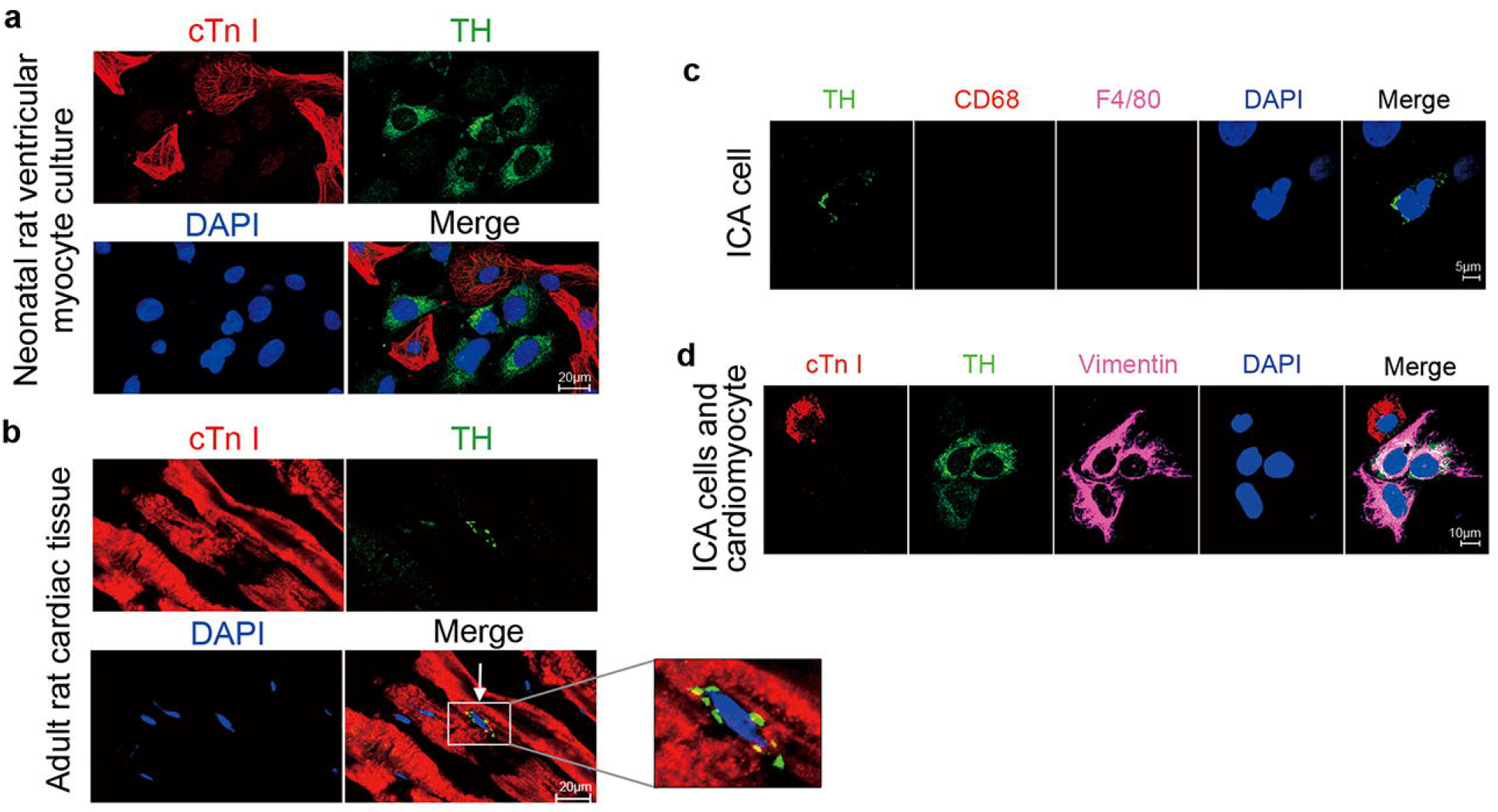
Intrinsic Cardiac Adrenergic (ICA) cells exist in neonatal and adult rat hearts. **a** and **b** Primary cultured neonatal rat cardiomyocytes isolated using traditional method and adult rat cardiac tissue. cTn I (cardiac troponin I): cardiomyocytes, red; TH (tyrosine hydroxylase): ICA cells, green. Cells are from *n*=6 neonatal rats in (**a**) and slides are from *n*=3 adult rats in (**b**). **c** ICA cells identified using macrophage markers, TH: ICA cell, green; CD68: marker of macrophage, red; F4/80: marker of macrophage, magenta. Cells are from *n*=6 neonatal rats. **d** ICA cells and cardiomyocyte identified using vimentin, cTn I: cardiomyocyte, red; TH: ICA cells, green; Vimentin: marker of mesenchymal cells, magenta. Cells are from *n*=6 neonatal rats. Nuclei of cells in all groups were dyed using DAPI: blue.

### ICA cells increase NE release that promotes cardiomyocyte TNF-α production in response to LPS stimulation

The primary NRVM culture containing ICA cells was defined as ICA cell-NRVM co-culture (NRVM ^ICA+^). To analyze the role of ICA cells, we used superparamagnetic iron oxide particles (SIOP) to remove ICA cells to obtain pure NRVM without ICA cell (NRVM ^ICA-^) (Fig. 2a).^24^ LPS stimulation significantly raised NE levels in the supernatants of NRVM ^ICA+^ in both a dose-dependent (LPS 0.01 μg/mL, 0.1 μg/mL, and 1 μg/mL) and time-dependent (LPS 0 h, 6 h and 12 h) manner when compared with controls (Fig. 2b). During the Langendorff perfusion of adult rat hearts, LPS elicited negligible influence on NE levels in perfusate, suggesting that circulating NE was not involved in this case due to the lack of sympathetic nerves and adrenal medulla (Fig. 2c). But, LPS administration significantly increased the level of NE in myocardial homogenate (Fig. 2d). Due to the presence of ICA cells in the NRVM ^ICA^ ^cell+^ culture and adult rat hearts, the increase of NE in the culture medium and myocardial homogenates could be attributed to ICA cells. To this end, we next examined and found that blockade of β_1_-AR using CGP20712A, a selective β_1_-AR antagonist, dramatically suppressed LPS-induced TNF-α production in NRVM ^ICA+^ culture, while removal of ICA cells abolished the impact of CGP20712A on TNF-α production by NRVM ^ICA-^ (Fig. 2e). Simultaneously, removal of ICA cells caused a lower level of TNF-α in the supernatants of LPS-stimulated NRVM ^ICA-^ than that of NRVM ^ICA+^ (Fig. 2f). These data support the concept that LPS stimulates ICA cell-derived NE release, which promotes LPS-induced TNF-α production by NRVM.

**Fig. 2.**
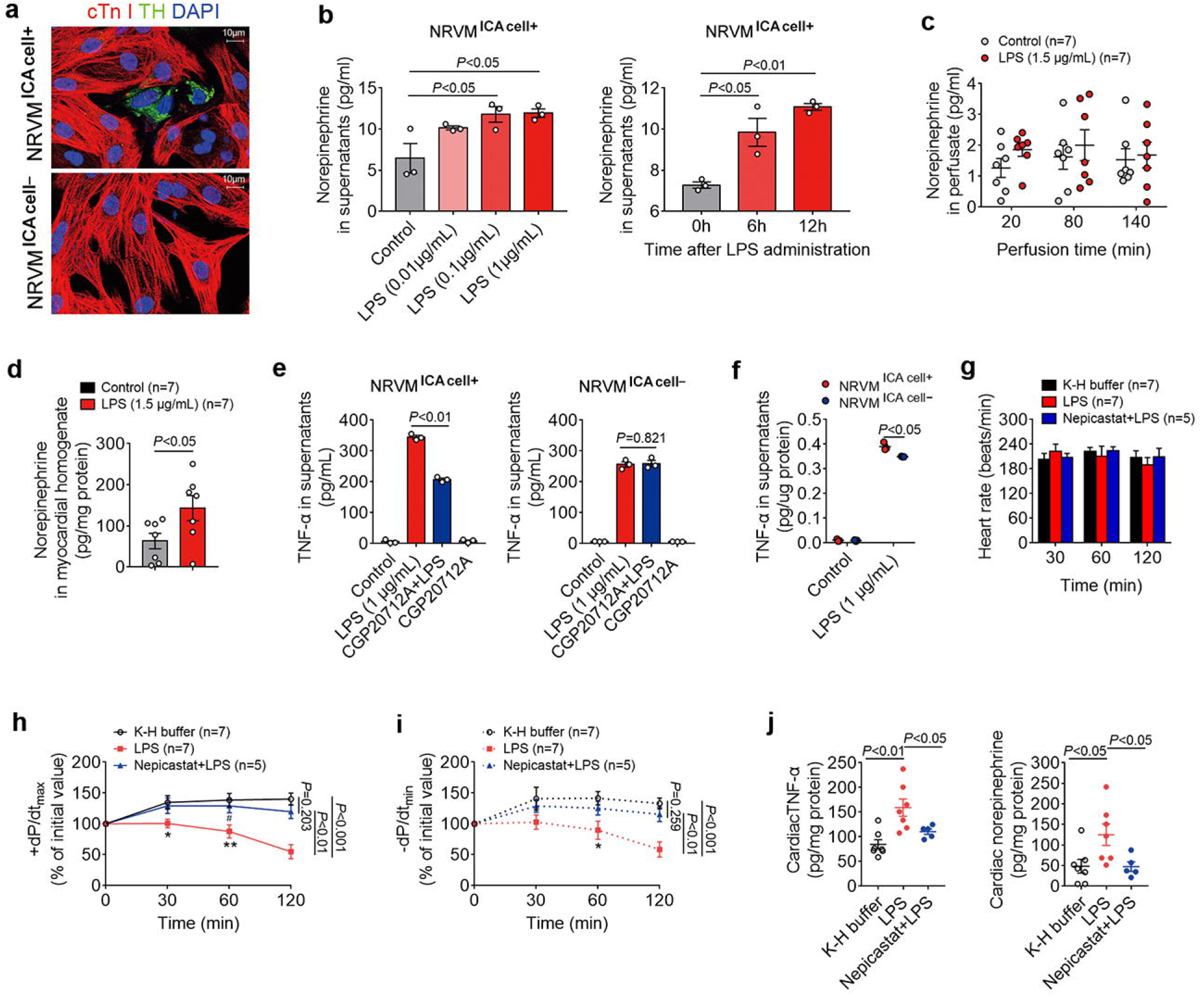
ICA cell-derived NE increased by LPS enhances LPS-induced cardiomyocyte TNF-α production and myocardial dysfunction. **a** ICA cell-neonatal rat ventricular myocyte co-culture (NRVM ^ICA+^) and NRVM without ICA cells (NRVM ^ICA-^), cTn I: cardiomyocytes, red; TH: ICA cells, green; DAPI: nuclei, blue. Cells are from *n*=12 neonatal rats. **b** Norepinephrine (NE) concentration in supernatants of NRVM ^ICA+^ stimulated with different doses of LPS for 6 h, and with 1 µg/mL LPS for different time points, *n*=3 independent experiments, cell density=5.5*10^5^ cells/mL. **c** NE in perfusate from Langendorff perfused adult rat hearts, control: K-H buffer. **d** NE in myocardial homogenates from adult rat hearts Langendorff perfused for 140 min, control: K-H buffer. Data are from *n*=7 rats (Control), *n*=7 rats (LPS group) in (**c** and **d**). **e** TNF-α in supernatants of NRVM ^ICA+^ and NRVM ^ICA-^ treated with CGP20712A, a β_1_-adrenergic receptor (AR) blocker, 2µM for 30 min prior to LPS for 6 h. **f** TNF-α in supernatants of NRVM ^ICA+^ and NRVM ^ICA-^ stimulated with 1 µg/mL LPS for 6 h. *n*=3 independent cell experiments, cell density=5.5*10^5^ cells/mL in (**e** and **f**). **g**-**i** Heart rate, systolic (+dP/dt_max_) and diastolic (-dP/dt_min_) function of Langendorff perfused adult rat hearts, LPS: 1.5 µg/mL, Nepicastat: Dopamine-β-hydroxylase (DBH) inhibitor, 15 µg/mL. **j** TNF-α and NE in myocardial homogenates from adult rat hearts Langendorff perfused for 120 min. Data in (**g**-**j**) are from *n*=7 rats (K-H buffer), *n*=7 rats (1.5 µg/mL LPS), *n*=5 rats (15 µg/mL Nepicastat+LPS). Data are presented as mean ± S.E.M. and analyzed using one-way ANOVA with Bonferroni post hoc test or two-tailed independent Student’s t-test as appropriate, \**P*<0.05, \*\**P*<0.01, most of the *P*-values were assigned on the figure. Independent cell experiments are biological replicates.

### Blockade of ICA cell-derived NE biosynthesis reduces LPS-induced cardiac TNF-α production and prevents myocardial dysfunction

To further investigate the function of ICA cells in myocardial dysfunction induced by LPS, we set up isolated adult rat hearts in the Langendorff perfusion system to remove the impact of sympathetic nerve-derived and adrenal medulla-sourced catecholamines as described previously.^26^ The Langendorff-perfused hearts in different groups showed a negligible alteration in heart rate (Fig. 2g). But, rat hearts in the LPS group exhibited a significantly worse phenotype in both systolic (+dP/dt_max_) and diastolic (-dP/dt_min_) functions compared with the K-H buffer group (control). By contrast, blockade of ICA cell-derived NE biosynthesis using Nepicastat, an inhibitor of DBH, resulted in a marked reversal in both systolic and diastolic dysfunctions compared with the LPS group (Fig. 2h, i). Nepicastat simultaneously abrogated the elevation of NE and TNF-α production in LPS-perfused hearts, consistent with changes in systolic and diastolic functions (Fig. 2j). These findings demonstrate that blockade of ICA cell-derived NE biosynthesis reduces LPS-induced cardiac TNF-α production and protects against LPS-induced myocardial dysfunction.

### NE-producing enzyme expression in ICA cells is upregulated by LPS stimulation

Given that TH and DBH are the key enzymes responsible for NE biosynthesis,^9^ we therefore evaluated the TH and DBH expression in ICA cells. Immuno-staining showed that LPS significantly stimulated TH expression in neonatal rat ICA cells as well as in Langendorff-perfused adult rat heart ICA cells, but not cardiomyocytes (Fig. 3a, b). In NRVM ^ICA+^, LPS (1μg/mL) stimulation for 6 h and 12 h dramatically increased TH and DBH mRNA expression, and different doses of LPS also significantly enhanced TH and DBH mRNA expression after 6 h stimulation compared to controls (Fig. 3c). Consistent with the upregulation in mRNA level, LPS markedly increased TH and DBH protein production in both a time- and dose-dependent manner as well (Fig. 3d, e). Similarly, in Langendorff-perfused adult rat hearts, LPS significantly increased TH and DBH expression in both of mRNA and protein levels compared to controls (Fig. 3f, g). These data show that LPS increase TH and DBH expression in ICA cells.

**Fig. 3.**
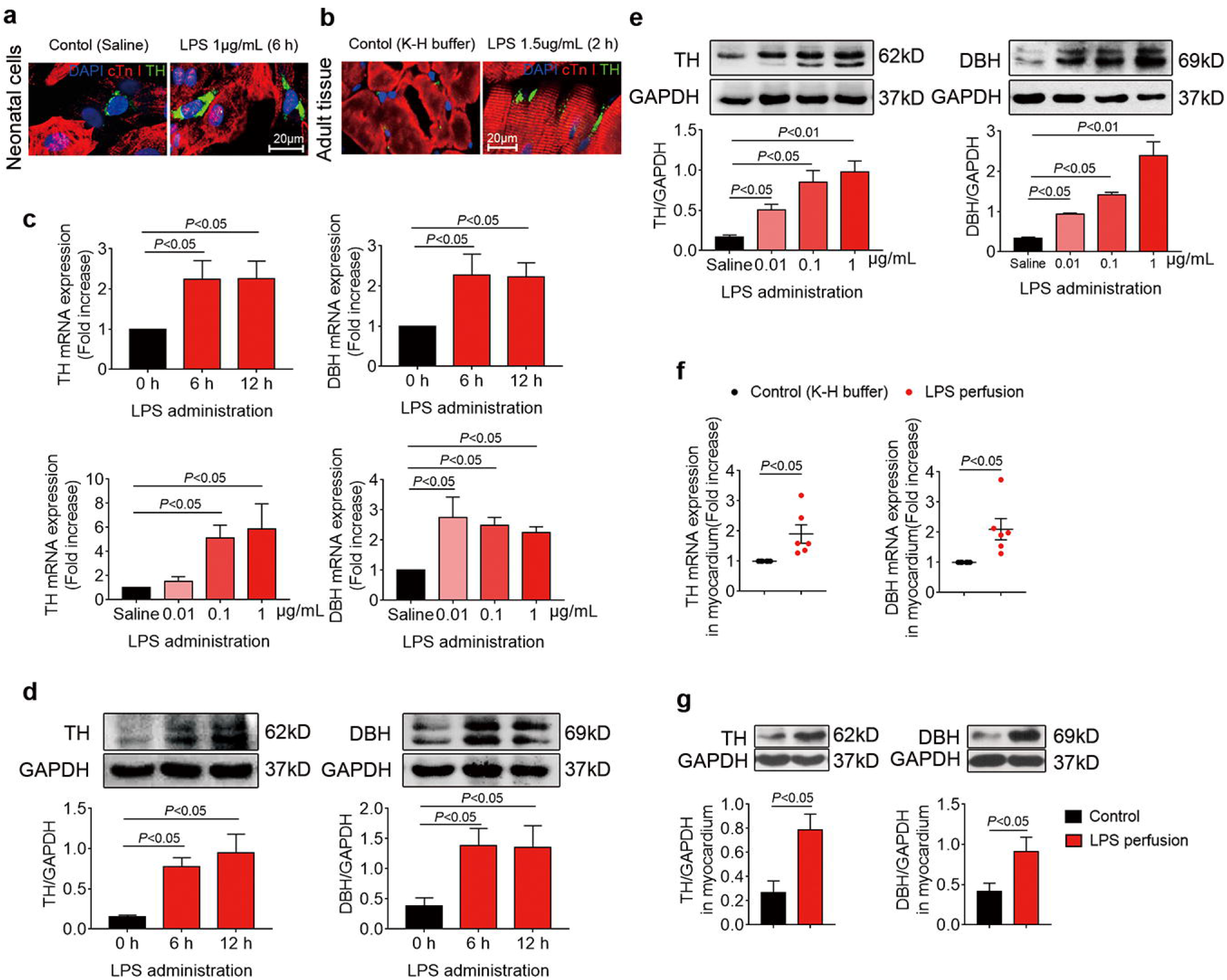
Expression of NE-producing enzymes in ICA cells is upregulated with LPS stimulation. **a** and **b** TH expression in NRVM ^ICA+^ treated with saline or LPS for 6 h and adult rat hearts Lagendorff perfused for 2 h; cTn I: cardiomyocytes, red; TH: ICA cells, green; DAPI: nuclei, blue. Cells are from *n*=6 neonatal rats in (**a**). Slides are from *n*=6 rats (Control), *n*=6 rats (LPS group) in (**b**). **c**-**e** mRNA and protein expression of TH and DBH relative to GAPDH in NRVM ^ICA+^ stimulated with LPS at different time points and doses. *n*=3 independent experiments in qPCR and Western blot, respectively, cell density=5.5*10^5^ cells/mL. **f** and **g** mRNA and protein expression of TH and DBH relative to GAPDH in myocardium from adult rat hearts Lagendorff perfused for 2 h, control (K-H buffer): *n*=6 rats, LPS (1.5 µg/mL in K-H buffer): *n*=6 rats. Data are presented as mean ± S.E.M. and analyzed using one-way ANOVA with Bonferroni post hoc test or two-tailed independent Student’s t-test as appropriate, *P*-values were assigned on the figure. Independent cell experiments are biological replicates.

### TLR4-MyD88/TRIF-AP-1 signaling mediates LPS-stimulated NE-producing enzyme upregulation

We next attempted to pinpoint the underlying mechanism of NE-producing enzyme upregulation in ICA cells stimulated by LPS. Since TLR4 is a well-known receptor of LPS,^27^ it might also be hypothesized that TLR4 signaling mediates LPS-induced upregulation of TH and DBH in ICA cells. We therefore examined the expression of TLR4 in ICA cells. ICA cells were isolated using the SIOP method (Fig. S2). Immuno-staining of TLR4 in NRVM^ICA^ ^cell+^ and isolated ICA cells found that ICA cells expressed TLR4, same as cardiomyocytes (Fig. 4a and Fig. S3A). Blockade of TLR4 signaling using Viper peptide (a blocker of TLR4) ^28^ markedly suppressed LPS-induced NE production as well as expression of TH and DBH in ICA cells (Fig. 4b, c). Consistently, DBH expression and NE production were not augmented in LPS-treated TLR4-deficient (*Tlr4*^*Lps-del*^) NMVM^ICA^ ^cell+^ (co-culture of ICA cells and neonatal mouse ventricular myocytes), which were extremely lower than those in LPS-treated wild-type controls (Fig. 4d-f). Furthermore, we cloned the rat TH promoter luciferase reporter (Fig. S3B-F), and found that LPS stimulation markedly activated the TH promoter as well as the AP-1 (Fig. 4g). Similarly, overexpression of the TLR4 adaptors, MyD88 and TRIF, significantly activate the TH promoter and the AP-1 as well (Fig. 4h and Fig. S3H). Given that c-Jun, as a downstream signal of TLR4 pathway, has specific binding sites on TH promoter,^29^ we therefore used siRNAs to knock down c-Jun and c-Fos protein levels to investigate the impact of AP-1 binding deficiency on TH promoter activation (Fig. 4i, Fig. S4). The activation of the TH promoter in response to MyD88 overexpression was dramatically suppressed by c-Jun siRNA and c-Fos siRNA (Fig. 4j). Single-cell RNA sequencing also verified that LPS induced the augmented trend of *Jun* expression in ICA cells (Fig. 5h). These results uncover that LPS stimulates NE-producing enzyme biosynthesis via TLR4-MyD88/TRIF-AP1 signaling activating TH promoter.

**Fig. 4.**
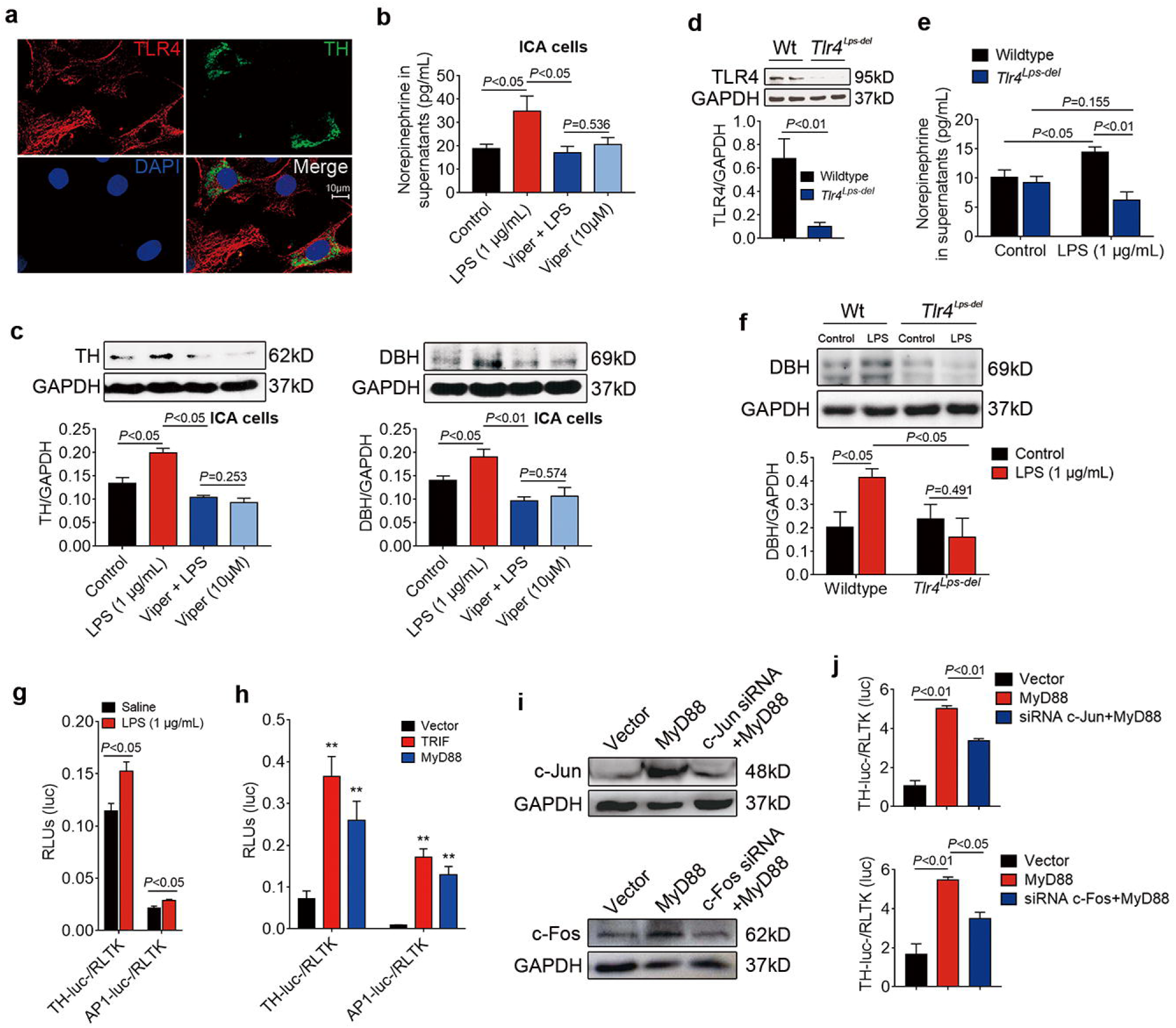
TLR4/AP-1 signaling mediates LPS upregulation of NE-producing enzyme expression. **a** Co-immunofluorescent staining of TLR4 and TH in NRVM ^ICA+^, TLR4: toll-like receptor 4, red; TH: ICA cells, green; DAPI: nuclei, blue. Cells are from *n*=6 neonatal rats. **b** and **c** NE production as well as TH and DBH expression in rat ICA cells treated with Viper (TLR4 inhibitor, 10 μM) for 2 h prior to LPS for 6 h. *n*=3 independent experiments, cell density=4.0*10^5^ cells/mL. **d** TLR4 expression in wildtype and TLR4-deficient (*Tlr4*^*Lps-del*^) NMVM^ICA^ ^cell+^. **e** and **f** NE production and DBH expression in wild-type and *Tlr4*^*Lps-del*^ NMVM^ICA^ ^cell+^ stimulated with LPS for 6h, control: saline. *n*=3 independent experiments, cell density=5.0*10^5^ cells/mL in (**d**-**f**). **g** Luciferase activity of AP-1-luc- and TH-promoter-luc-relative to RLTK-luc-in Hek293/hTLR4 cells at 36 h after transfection and LPS stimulation. **h** Luciferase activity of AP-1-luc- and TH-promoter-luc-relative to RLTK-luc-in Hek293/hTLR4 cells at 24 h after transfection with plasmids (0.1 μg/mL) encoding MyD88 and TRIF. **i** Expression of c-Jun and c-Fos in 293/hTLR4 cells at 24 h after transfection with MyD88 plasmid, c-Jun siRNA and c-Fos siRNA. **j** Luciferase activity of TH-promoter-luc-relative to RLTK-luc-in Hek293/hTLR4 cells at 24 h after transfection with MyD88 plasmid, c-Jun siRNA and c-Fos siRNA, (0.1 μg/mL for plasmid). *n*=3 independent experiments, cell density=4.5*10^5^ cells/mL in (**g**-**j**). Data are presented as mean ± S.E.M. and analyzed using one-way ANOVA with Bonferroni post hoc test or two-tailed independent Student’s t-test as appropriate, \**P*<0.05, \*\**P*<0.01, most of the *P*-values were assigned on the figure. Independent cell experiments are biological replicates.

**Fig. 5.**
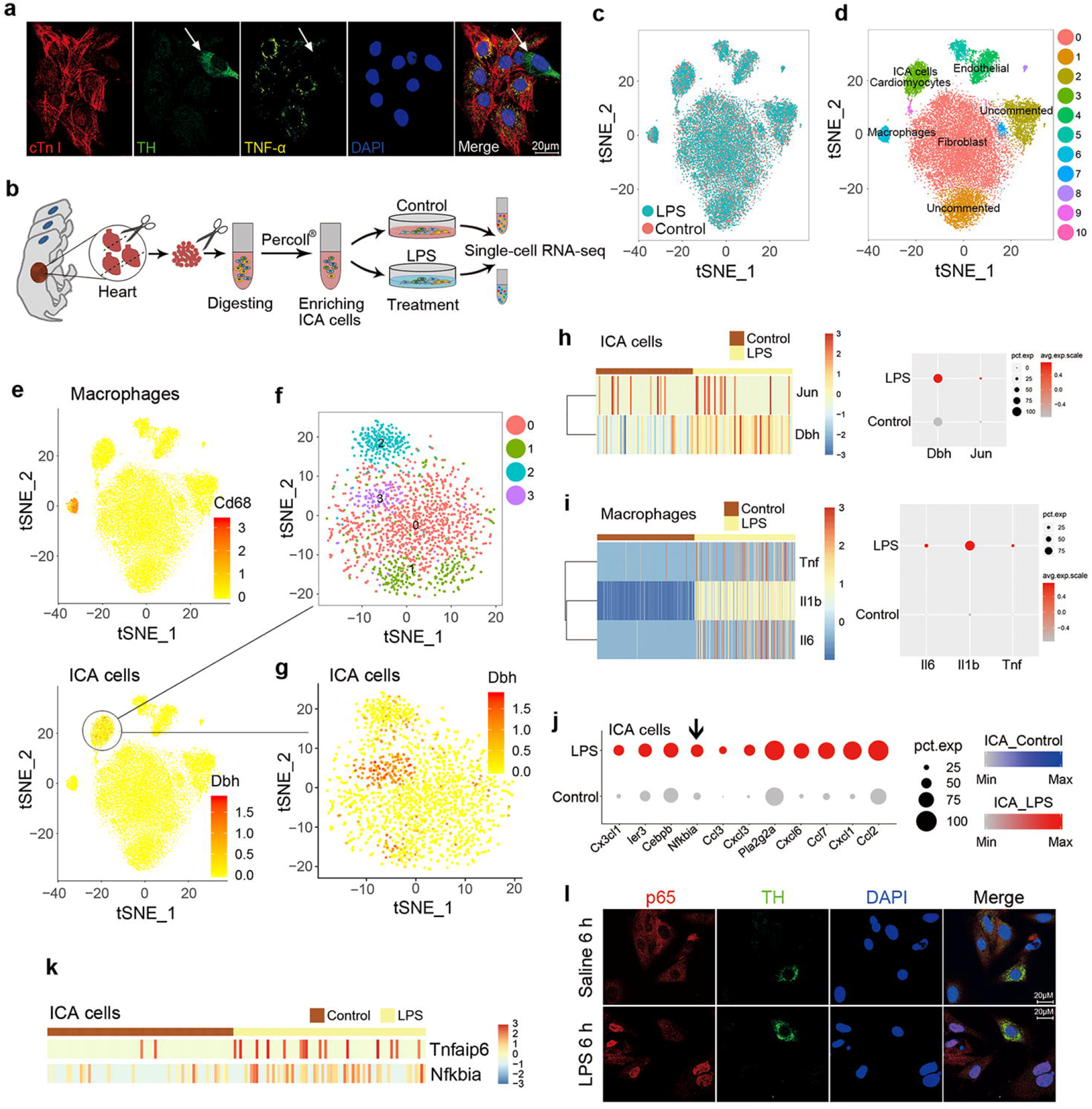
ICA cell does not produce TNF-α with LPS stimulation. **a** Immunofluorescent-staining of TNF-α in NRVM ^ICA+^ stimulated with 1 µg/mL LPS for 6 h, cTn I: cardiomyocytes, red; TH: ICA cells, green; TNF-α, yellow; DAPI: nuclei, blue. Cells are from *n*=12 neonatal rats. **b** Schematic of single-cell RNA-sequencing experimental strategy. **c** *t*-distributed stochastic neighbor embedding (t-SNE) projection of cardiac cells, batch effect analysis between control and LPS groups, control: saline, *n*=24210 cells. **d** t-SNE map of cardiac cells, different colored clusters represent distinct populations, *n*=24210 cells. **e** ICA cells expressing *Dbh* and macrophages expressing *Cd68* are separated in two clusters, *n*=24210 cells. **f** and **g** Analysis of substructure in ICA and cardiomyocyte cluster indicates subcluster 3 (Purple) is ICA cell subpopulation, *n*=1765 cells. **h** Expression of *Dbh* and *Jun* in ICA cells between control and LPS groups, no *Tnf* gene expression, *n*=138 cells. **i** Expression of *Tnf, Il6* and *Il1b* in macrophages between control and LPS groups, *n*=572 cells. **j** Top11 of variable differentially expressed genes (DEGs) which include *Nfkbia* in the subcluster ICA cells between control and LPS groups, control: saline, *n*=138 cells. **k** Expression of *Tnfaip6* and *Nfkbia* in ICA cells between control and LPS groups, *n*=138 cells. **l** Immunofluorescent-staining of p65 localization in NRVM ^ICA+^ stimulated with saline or 1 µg/mL LPS for 6 h, p65, red; TH: ICA cells, green; DAPI: nuclei, blue. Cells are from *n*=6 neonatal rats. For single-cell transcriptomes, cells are from 24 neonatal rats. Gene-expression values represent mean of log; differential expression test: Wilcoxon rank sum test; avg_logFC= log (mean(group1)/mean(group2)); adjusted p_value: Bonferroni Correction; min.pct ≥10%; avg_logFC >= 0.1.

### TLR4-expressing ICA cell does not produce TNF-α with LPS stimulation

In order to further clarify the role of ICA cells in LPS-induced cardiac TNF-α production, we need to know whether ICA cells possess the capability of TNF-α production. To this end, we performed co-immunofluorescence staining of TNF-α, TH and cTn I in LPS sitmulated NRVM ^ICA+^, and found that TNF-α was definitely induced in cardiomyocytes but not in ICA cells by LPS (Fig. 5a). Subsequently, single-cell RNA sequencing was utilized to validate the phenotype and pinpoint the mechanisms. Any batch-from-batch variations between the control and LPS groups were removed by normalizing the two-groups of data (Fig. 5c). Distinct populations of cardiac cells were identified in different clusters on *t*-SNE map (Fig. 5d). Unsupervised clustering revealed macrophages expressing *Cd68* and ICA cells expressing *Dbh* in separate clusters, supporting our immuno-staining results that ICA cells were not macrophages (Fig. 5d, e). Analysis of substructure in cluster 3 (ICA cells and cardiomyocytes) showed four distinct groups in which the new sub-cluster 3 (purple) was the right population of ICA cells expressing *Dbh* (Fig. 5f, g). Close-up of the differentially expressed genes (DEGs) heat maps and dot plots displayed an upregulation of *Dbh* and *Jun* expression, but no *Tnf* expression in LPS-treated ICA cells (Fig. 5h). While as a positive control, *Tnf, Il1b* and *Il6* gene expression were all upregulated in LPS-stimulated macrophages (Fig. 5i). Mechanistically, analyses of DEGs showed significantly increased expressions of *Nfkbia* and *Tnfaip6* in LPS-treated ICA cells compared with controls (Fig. 5j,k and Fig. S5C, D). Since *Nfkbia* is involved in p65 nuclear translocation,^30^ and TNF-α-stimulated gene-6 (TSG6) coded by the gene *Tnfaip6* also suppresses NF-κB activation,^31^ the upregulation of *Nfkbia* and *Tnfaip6* could be the reason of the lack of the capability of TNF-α production in ICA cells. We therefore performed immunofluorescence staining of p65 localization to examine NF-κB activation in LPS-treated NRVM ^ICA+^. As expected, LPS stimulation did not activate the nuclear translocation of p65 in ICA cells, while significant nuclear translocation of p65 was found in LPS-treated non-ICA cells compared to saline-treated controls (Fig. 5l). These findings demonstrate that ICA cell loses the capacity of TNF-α production due to the inhibition of p65 nuclear translocation by the elevation of *Nfkbia* and *Tnfaip6* with LPS stimulation.

### ICA cell-derived NE acts via β_1_-AR-CaMKII signaling to regulate NF-κB and MAPK pathways in cardiomyocyte

To understand how ICA cell-derived NE interacts with proinflammatory signaling in NRVM, effects of different adrenergic receptor blockade were observed, where only β_1_-AR or β_2_-AR blockade significantly reduced the LPS-induced TNF-α production in NRVM ^ICA+^ (Fig. S6A). Administration of β_1_-AR blocker, CGP20712A (2 µM), prior to LPS (1 μg/mL) treatment dramatically suppressed LPS-induced phosphorylation of IκBa and p65 (Fig. 6a), which fits well with previous work that activation of β_1_-AR increases IκBα phosphorylation and TNF-α expression in septic mouse hearts.^11^ In addition, the levels of phosphorylated-JNK and phosphorylated-p38 in the CGP20712A+LPS group were significantly decreased, while the expression of c-Fos and phosphorylated-ERK1/2 was evidently increased compared to the LPS group (Fig. 6b, c). These data are supported by other reports that ERK1/2 phosphorylation raises c-Fos expression, subsequently inhibiting p38 MAPK phosphorylation to reduce cardiomyocyte TNF-α production in endotoxemia.^22,32^ These results suggest that the NF-κB and MAPK signaling pathways were involved in the process of ICA cell-derived NE promotion of LPS-induced cardiomyocyte TNF-α production. Since protein kinase A (PKA) signaling has proven to be a major route for channeling the cardiac β_1_-AR signaling,^33^ we tested the phosphorylation of CREB-Ser133 which indicates PKA activation.^34^ Intriguingly, PKA activation showed no significant alteration in all of the groups (Fig. 6d). Yet, the phosphorylation of CaMKII was dramatically enhanced in LPS-stimulated NRVM ^ICA+^ compared with controls, while CGP20712A significantly reduced CaMKII phosphorylation compared to LPS group (Fig. 6e). These data suggested that CaMKII phosphorylation is a critical signaling involving in this progress. We next used KN93, a selective inhibitor of CaMKII, to block CaMKII signaling. KN93 markedly decreased the levels of TNF-α in the supernatants of LPS-treated NRVM ^ICA+^ in a dose-dependent manner (Fig. 6f). The phosphorylation of CaMKII in the KN93+LPS group was also significantly reduced compared with LPS-treated NRVM ^ICA+^ (Fig. 6g). Accordingly, KN93 dramatically decreased LPS-induced phosphorylation of p65, IkBa, p38 and JNK in NRVM ^ICA+^ (Fig. 6h, i), and increased ERK1/2 phosphorylation (Fig, 6j). These results show that CaMKII blockade elicits a similar phenotype as what β_1_-AR blockade does in LPS-treated NRVM ^ICA+^, which suggests that CaMKII is essential during the β_1_-AR signal transduction process activated by ICA cell-derived NE.

**Fig. 6.**
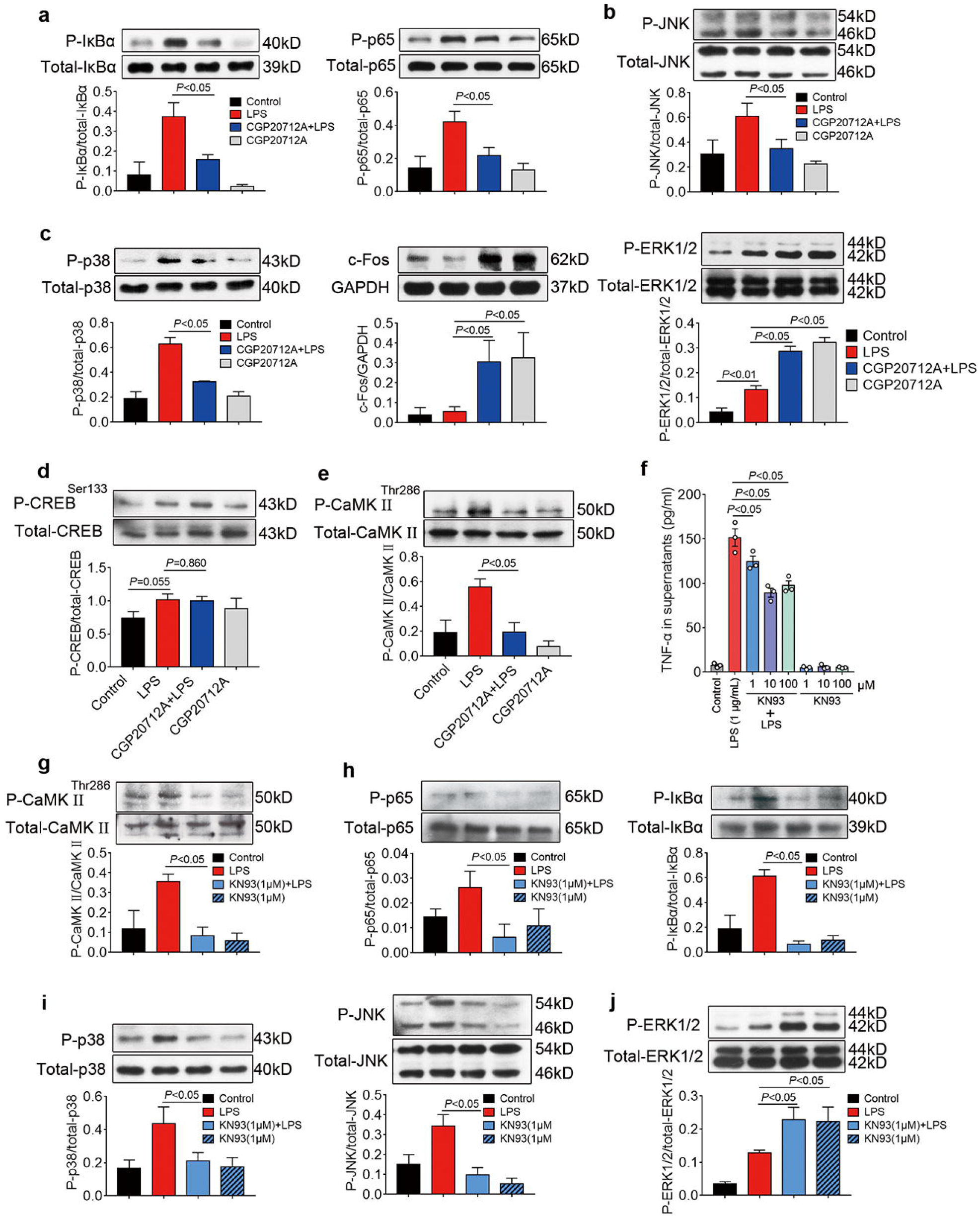
ICA cell-derived NE enhanced by LPS acts via β_1_-AR-CaMKII pathway to regulate NF-κB and MAPK signaling pathways. **a**-**e** NRVM ^ICA+^ were stimulated with 2 µM CGP20712A (β_1_-AR antagonist) for 30 min prior to 1 µg/mL LPS for 30 min, (**a**) phosphorylation of IκBα and p65, (**b**) phosphorylation of JNK, (**c**) phosphorylation of p38, expression of c-Fos and ERK1/2 phosphorylation, (**d**) phosphorylation of CREB-Ser133, indicating PKA activation. (**e)** phosphorylation of CaMKII in NRVM ^ICA+^. **f** TNF-α in supernatants of NRVM ^ICA+^ treated with KN93 in different doses for 30 min prior to 1 µg/mL LPS for 6 h. **g-j** Phosphorylation of CaMKII, p65, IκBα, p38, JNK and ERK1/2 in NRVM ^ICA+^ treated with 1 µM KN93 for 30 min prior to 1 µg/mL LPS for 30min. For all of the cell experiments, *n*=3 independent experiments, cell density=5.5*10^5^ cells/mL. Data are presented as mean ± S.E.M. and analyzed using one-way ANOVA with Bonferroni post hoc test or two-tailed independent Student’s t-test as appropriate, *P*-values were assigned on the figure. Independent cell experiments are biological replicates.

### Dobutamine stimulation on β_1_-AR recapitulates the impact of ICA cell-derived NE on TNF-α production in LPS-treated cardiomyocyte

In order to validate our findings, dobutamine (DOB), a selective β_1_-AR agonist,^35^ was employed to mimic ICA cell-derived NE to treat NRVM without ICA cells (NRVM ^ICA-^) prior to LPS stimulation. We found that short-term (10-minute) β_1_-AR stimulation with DOB sharply reduced the LPS-induced TNF-α production in NRVM ^ICA-^, while PKA inhibitor 14-22 amide (PKI) increased LPS-induced TNF-α production and reversed the suppressed effect of DOB (Fig. 7a). Such observations fit well with published data that short-term β_1_-adrenergic stimulation elicited a rapid increase of cellular cAMP activating PKA,^36^ which inhibited LPS-induced cardiac expression of TNF-α.^37^ Of note, cardiomyocytes are regularly, but not short-time, regulated by catecholamine due to the physiological intrinsic cardiac adrenergic activities. We thus prolong the treatment time of DOB, and found that β_1_-AR stimulation with DOB for 6 h significantly increased the level of TNF-α in supernatants of LPS-treated NRVM ^ICA-^ (Fig. 7b). Same as ICA cell-derived NE, DOB stimulation of NRVM ^ICA-^ for 6 h did not alter phosphorylation of CREB-Ser133 (Fig. 7c), but dramatically enhanced the level of phosphorylated-CaMKII^Thr286^ compared with control and LPS groups (Fig. 7d). The phosphorylation of CaMKII^Thr286^ in LPS-treated NRVM ^ICA-^ showed negligible differences compared with controls, indicating that there was no NE production due to the lack of ICA cells in NRVM ^ICA-^ culture (Fig. 7d). Consistent with the results of β_1_-AR blockade and CaMKII blockade in LPS-stimulated NRVM and ICA cell co-culture (NRVM ^ICA+^), DOB stimulation of NRVM ^ICA-^ for 6 h markedly increased phosphorylation of IκBα and JNK and decreased the level of phosphorylated-ERK1/2 compared with LPS-treated NRVM ^ICA-^, whereas no change in phosphorylation of p38 in NRVM ^ICA-^ (Fig. 7e, f). The unchanged p38 phosphorylation may because of abolished effect of the mild β_2_-AR agonist activity of DOB.^38^ It has been demonstrated that β_2_-AR activation suppressed p38 MAPK phosphorylation.^39^ These data suggest that DOB promoted TNF-α production in LPS-stimulated NRVM, which successfully mimicked ICA-cell derived NE, and therefore validated our findings that ICA-cell derived NE contributes to LPS-induced TNF-α production via CaMKII-dependent β_1_-AR signaling crosstalk with NF-κB and MAPK signaling pathways in cardiomyocytes.

**Fig. 7.**
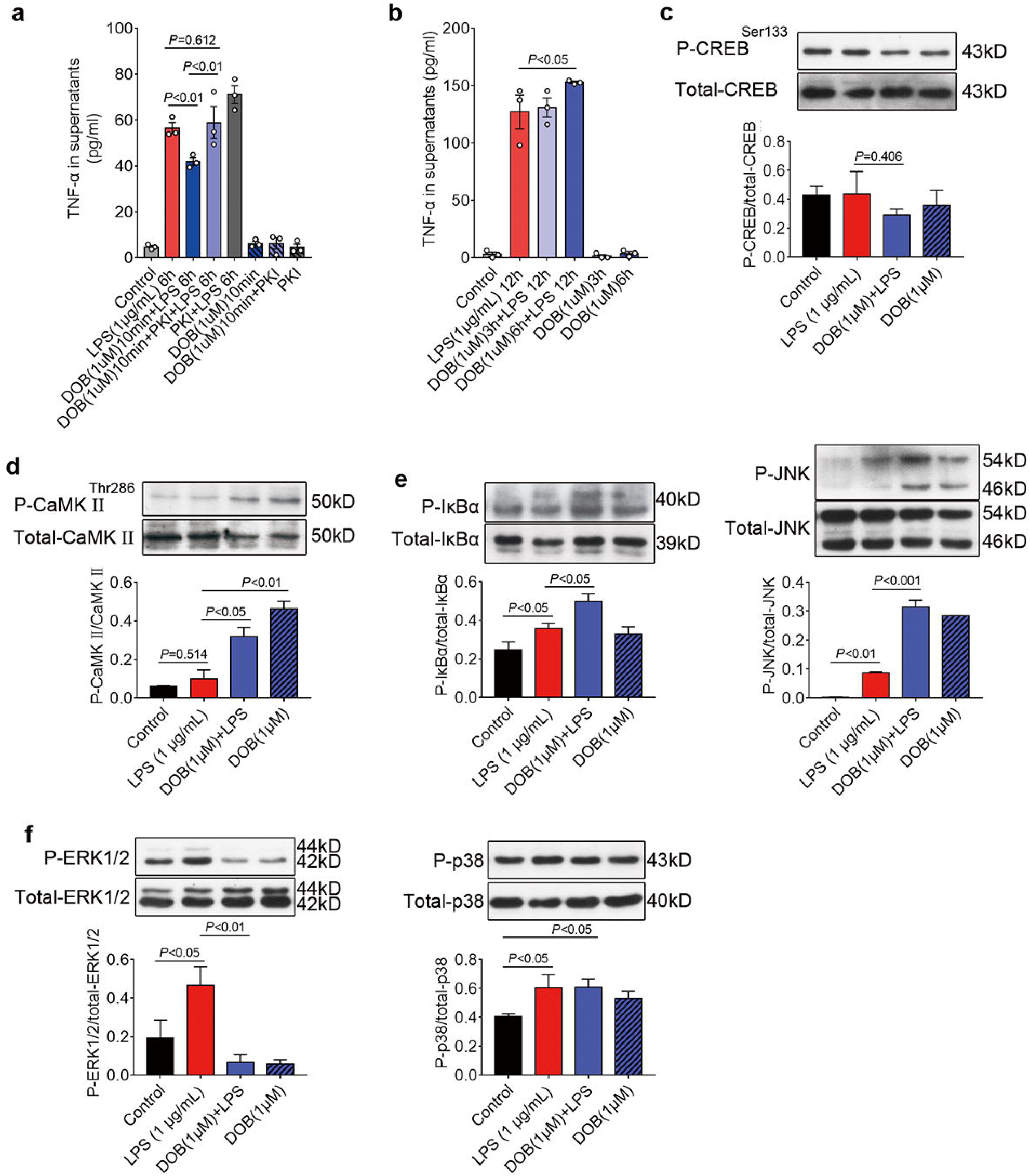
Dobutamine stimulation on β_**1**_**-AR recapitulates the impacts of ICA cell-derived NE on TNF-**α **production in LPS treated NRVM** ^**ICA-**^. **a** TNF-α in supernatants of NRVM without ICA cell (NRVM ^ICA-^) treated with 1 µM dobutamine (DOB) for 10 min and/or 5 µM PKA inhibitor 14-22 amide (PKI) for 1 h prior to LPS 1 µg/mL for 6 h. **b** TNF-α in supernatants of NRVM ^ICA-^ treated with 1 µM DOB for 3 h or 6 h prior to 1 µg/mL LPS for 12 h. **c-f** Phosphorylation of CREB-Ser133, CaMKII-Thr286, IκBα, JNK, ERK1/2 and p38 in NRVM ^ICA-^ treated with 1 µM DOB for 6 h prior to 1 µg/mL LPS for 30min. For all of the cell experiments, *n*=3 independent experiments, cell density=5.0*10^5^ cells/mL. Data are presented as mean ± S.E.M. and analyzed using one-way ANOVA with Bonferroni post hoc test or two-tailed independent Student’s t-test as appropriate, *P*-values were assigned on the figure. Independent cell experiments are biological replicates.

## Discussion

Catecholamines are irreplaceable in both of developing and adult hearts. Targeted disruption in mice of the genes encoding catecholamine biosynthesis enzymes are embryonic lethal, likely due to cardiac failure.^40,41^ The developing heart initially relies on ICA cells as the major source of catecholamines.^42^ Published evidences have shown that ICA cells synthesize cardiac intrinsic catecholamines occurring independently of cardiac sympathetic nerves either in neonatal or adult hearts, thereby functioning as a critical and integral regulator in mammalian heart development, cardiac pathophysiology and post-heart transplantation support.^14-17,19,21,43,44^ Stimulation of ICA cells enhances epinephrine release to reduce ischemia/reperfusion injury through δ-opioid-regulated cardioprotective adrenopeptidergic co-signaling pathway.^17,19,45^ Although ICA cell density shows negligible connection with reduced myocardial efficiency in failing myocardium,^18^ irregular stimulation of ICA cells increases catecholamine-synthesizing enzymes and cardiac norepinephrine levels.^46^ Such actions of ICA cells may represent an important contributive mechanism in septic hearts since increased cardiac norepinephrine and β_1_-AR stimulation aggravate sepsis-induced cardiomyocyte apoptosis and myocardial injury.^8,11^ However, to date the potentially important role of ICA cell in sepsis-induced cardiac dysfunction is rarely examined and thus remains largely unknown.

Indeed, ICA cells were not separated from cardiomyocytes in the most current cardiac researches using primary neonatal rat ventricular myocyte (NRVM) culture model to define function and attribute of cardiac cells.^47,48^ Our previous study used a superparamagnetic iron oxide particle (SIOP) method to obtain pure NRVM without ICA cells and demonstrated the ICA cell-free NRVM showed lower calcium transient amplitude, took longer to begin autonomous beating and expressed less natriuretic peptide A (Nppa) and -B (Nppb) (markers of stress response) in the condition of untreatment than those of NRVM mixed with ICA cells isolated by traditional methods.^24^ In addition, we also observed a markedly reduced LPS-induced TNF-α production in ICA cell-free NRVM compared to traditionally isolated NRVM which contains ICA cells during culture.^24^ Therefore, it is important to distinguish and evaluate the function of ICA cells in the cardiac study.

In this study, we defined ICA cell-derived NE as a contributor to facilitate myocardial proinflammatory response and dysfunction via activating β_1_-AR in cardiomyocytes during sepsis, and this was substantiated by the following evidences. Firstly, our results showed that ICA cells were not macrophages, but mesenchymal cells, and acted as a source of increased NE during LPS treatment in both traditionally cultured neonatal rat cardiomyocytes and adult rat hearts. NE levels were significantly raised in both the supernatants of NRVM ^ICA^ ^cell+^, ICA cells and the myocardial homogenate of Langendorff-perfused hearts after LPS challenge. Although endothelial cells have been evidenced to be able to produce catecholamines in response to ischemia,^49^ the typical “cobblestone” monolayer pattern of morphology in endothelial cell culture largely differs from ICA cells.^50^ The unchanged NE levels in the perfusate of LPS-perfused hearts also provided evidence that the cardiac NE level increased by LPS was not from vascular endothelial cells during Langendorff perfusion. Secondly, either removal of ICA cells or blockade of β_1_-AR markedly decreased LPS-induced TNF-α in NRVM ^ICA^ ^cell+^ culture, while β_1_-AR blockade exerted no influence on TNF-α production in ICA cell-free NRVM. Of note, ICA cells were found to be lack of production of TNF-α during LPS stimulation, evidenced by the negative immune-staining of cytoplasmic TNF-α and nuclear p65, as well as single-cell RNA sequencing of gene expression in ICA cells. Thirdly, β_1_-AR blockade decreased LPS-induced TNF-α production as well as phosphorylation of CaMKII, IκBα, p65, JNK and p38 in NRVM ^ICA^ ^cell+^, while β_1_-AR agonist mimicked ICA-cell derived NE bioactivity in LPS-stimulated ICA cell-free NRVM. Lastly, inhibition of ICA cell-derived NE synthesis by nepicastat, an inhibitor of DBH, prevented LPS-induced cardiac NE and TNF-α production as well as cardiac dysfunction of isolated rat hearts in the Langendorff system.

TLR4, mediating recognition of LPS, has been well-known as the first-line sentinel of host defense against bacteria in innate immunity.^27^ However, its function in the regulation of hormone or neurotransmitter remains poorly investigated. Prior study has shown that LPS continuously increases TH expression in mouse brains,^51^ but the underlying mechanism remains a mystery. In the present study, we discovered that TLR4 was present in ICA cells and its downstream MyD88/TRIF-AP-1 signaling pathway was responsible for LPS-induced catecholamine synthesis in these cells. LPS upregulated NE-producing enzyme, TH and DBH, expression at both gene and protein levels in ICA cells. Either TLR4 inhibition or knockout significantly suppressed LPS-induced TH and DBH expression as well as NE production in ICA cells. Moreover, both of TRIF and MyD88 activated TH-promoter, and either c-Jun siRNA or c-Fos siRNA markedly suppressed MyD88-induced TH-promoter activation, suggesting that AP-1 binding mediates TLR4-MyD88/TRIF-induced TH expression. These results demonstrated for the first time that TLR4/AP-1/TH signal pathway was responsible for LPS-induced biosynthesis of cellular catecholamine in ICA cells. The previous studies show that TLR4 plays a fundamental role in activation of host immunity and expressed on immune cells and neurons ^4,52^ which possess the property of catecholamine synthesis,^25,53^ our results identified that TLR4 signal pathway activation in ICA cells promoted catecholamine synthesis and cardiac dysfunction in the development of sepsis. This concept significantly improved the understanding of TLR4 functions.

Although the activation of TLR4 triggers the release of cytokines including TNF-α in cardiac cells,^27^ this is not the case in ICA cell that expresses TLR4. According to single-cell RNA sequencing, the *Nfkbia* and *Tnfaip6* expression was upregulated in LPS-treated ICA cells. Published evidence has demonstrated that degradation of IκBα, coded by *Nfkbia*, is necessary for activation of NF-κB (p65) nuclear translocation, which is one of the predominant mechanisms responsible for LPS-induced TNF-α production.^30^ A recent study also shows TSG6, coded by *Tnfaip6*, suppresses NF-κB activation via inhibiting the association of TLR4 with MyD88.^31^ Indeed, we found that exposure of ICA cells to LPS did not stimulate p65 nuclear translocation, whereas NRVM did. This is different from the well-established cognition of TLR4 function.^27^ These findings indicate that activation of TLR4 on ICA cells predominantly drives the synthesis of hormones, such as epinephrine and norepinephrine, rather than TNF-α production due to upregulated *Nfkbia* and *Tnfaip6* expression, during sepsis. However, the complicated effects of ICA cell-TLR4 activation on hormone regulation remain unclear, and further investigation is warranted to define the underlying mechanisms. In addition, the marked upregulation of chemokine family gene expression (i.e. *Cxc3cl1, ccl3, cxcl1, ccl2*, etc.) in LPS-treated ICA cells indicate that ICA cells may also contribute to cell migration involved in homeostatic and inflammatory processes.

Furthermore, we demonstrated that the TNF-α production and CaMKII phosphorylation in cardiomyocytes were dramatically augmented in LPS-treated NRVM ^ICA^ ^cell+^ as a consequence of ICA cell-derived NE elevation, which were reversed by β_1_-AR blockade. Inhibition of CaMKII recapitulated the impact of β_1_-AR blockade on TNF-α expression in LPS-stimulated NRVM ^ICA^ ^cell+^, as well as NF-κB and MAPK signaling activation, demonstrating that CaMKII is essential in this process. These findings are similar with the published evidence that sustained β_1_-adrenergic stimulation promotes myocyte apoptosis and modulates cardiac contractility by PKA-independent CaMKII signaling pathways in chronic heart failure.^36,54^ Intriguingly, although PKA activation was reported to be involved in the process of β_1_-AR promoting cardiomyocyte apoptosis in septic hearts,^11^ the present study found that short-term stimulation of β_1_-AR using dobutamine suppressed LPS-induced TNF-α expression in cardiomyocytes, while PKA inhibition reversed this suppressive efficacy, indicating that PKA is a negative regulator in this process. These results fit well with published data that short-term β_1_-adrenergic stimulation elicited a rapid increase of cellular cAMP activating PKA,^36^ which inhibited LPS-induced cardiac expression of TNF-α.^37^ However, the release of NE from ICA cells in NRVM ^ICA^ ^cell+^ culture is continuous,^23^ we found that CaMKII was not activated in LPS-treated NRVM without ICA cell, while DOB stimulation of β_1_-AR for 6 h significantly activated CaMKII, rather than PKA, and promoted LPS-induced TNF-α production by cardiomyocytes. Recent evidence showed that CaMKII oxidation in response to myocardial infarction contributed to NF-κB-dependent inflammatory transcription in cardiomyocytes.^55^ Another study also demonstrated that oxidation-activated CaMKII had a causal role in the contractile dysfunction during sepsis.^56^ Therefore, the present study demonstrated that ICA cell-derived NE was another factor in activating cardiomyocyte CaMKII via β_1_-AR to enhance LPS-induced cardiac inflammation, and the β_1_-AR-CaMKII signaling activation was the principal mechanism via which ICA cell-derived NE enhanced cardiomyocyte NF-κB and MAPK signaling to promote LPS-induced cardiomyocyte TNF-α production.

Taken together, our data revealed that LPS activated the promoter of tyrosine hydroxylase by AP-1 binding to promote NE biosynthesis in ICA cell that expresses TLR4, and NE derived from ICA cells served as a paracrine signal to aggravate LPS-induced myocardial TNF-α production and cardiac dysfunction via β_1_-AR-CaMKII signaling, which has been underestimated before. Strikingly, LPS-TLR4 signaling activation in ICA cells does not trigger TNF-α production, but rather promotes NE production, indicating a novel cell type-specific TLR4 function in regulating hormone release during sepsis (Fig. 8). Therefore, our findings identified a pivotal capacity of ICA cells in regulating LPS-induced myocardial dysfunction, ICA cell-derived NE may be potentially therapeutic targets in the fight against SIMD.

**Fig. 8.**
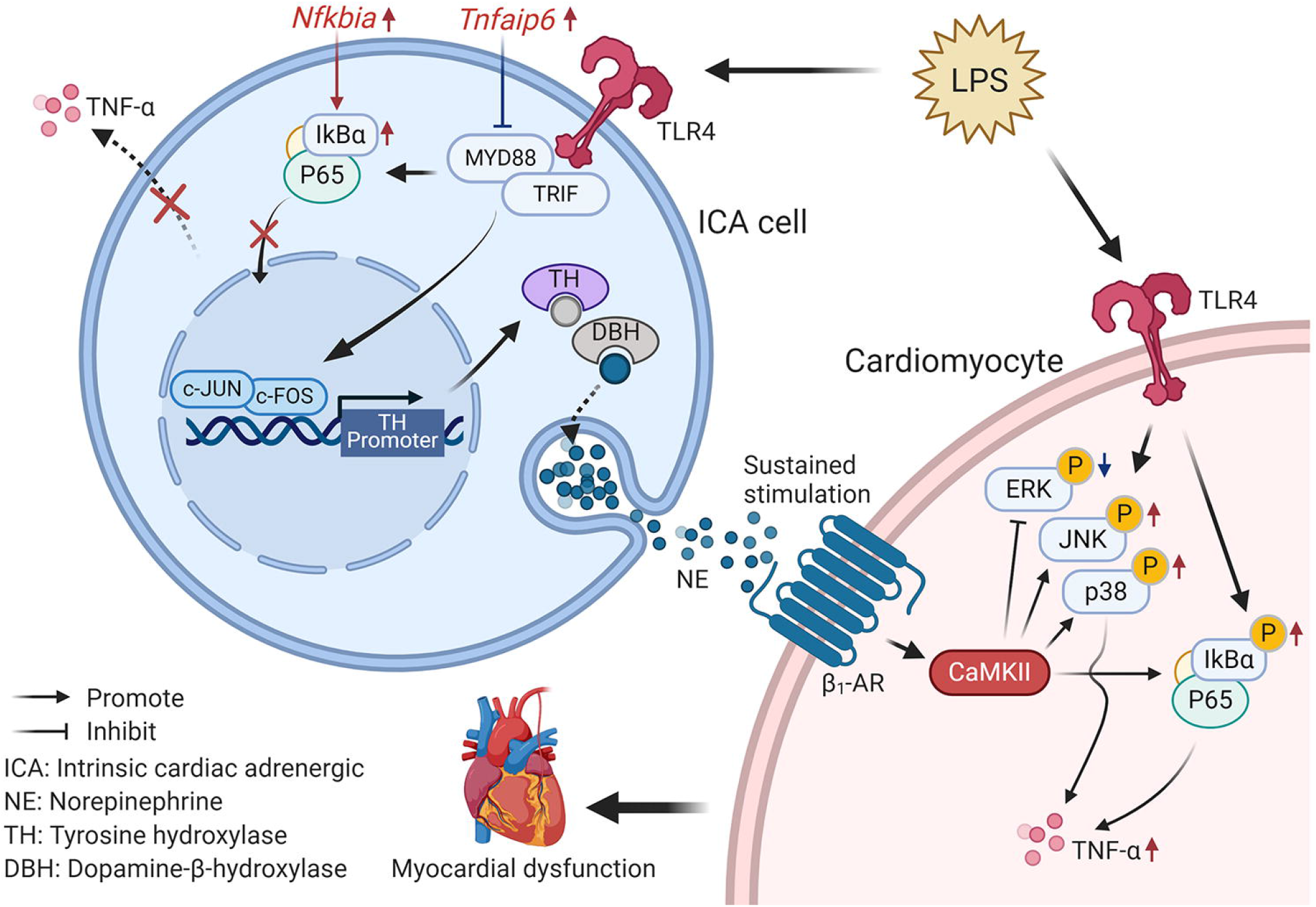
Schematic of ICA cell contribution to septic cardiomyopathy. ICA cell produces no TNF-α due to the elevated *Nfkbia* and *Tnfaip6* expression, but increases NE release by LPS-TLR4 signaling, which activates β1-AR-CaMKII signaling in cardiomyocyte to regulate NF-κB and mitogen-activated protein kinase pathways, subsequently aggravates myocardial TNF-α production and dysfunction.

## Materials and Methods

### Animals and cell line

All experiments with animals were conducted in compliance with the Guide for the Care and Use of Laboratory Animals published by the US National Institutes of Health ^57^ and approved by the Animal Care and Use Committee at Jinan University. Neonatal (1-3 days old) and adult (8-10 weeks) Sprague-Dawley rats were obtained from the laboratory animal center of Southern Medical University (Guangzhou, China). TLR4-deficient mice (*Tlr4*^*Lps-del*^, strain: C57BL/10ScNJNju) were purchased from Nanjing Biomedical Research Institute of Nanjing University. Hek293/hTLR4-HA cells (InvivoGen#293-htlr4ha) were a gift from Guang Yang (Jinan University).

### Inhibitors

α1-AR antagonist: Prazosin (Sigma-Aldrich#7791); α2-AR antagonist: Yohimbine (Sigma-Aldrich #Y3125); α2A-AR antagonist: BRL 44408 (Sigma-Aldrich #B4559); β1-AR antagonist: CGP20712A (Sigma-Aldrich #C231); β2-AR antagonist: ICI-118 551 (Sigma-Aldrich #I127); β3-AR antagonist: SR59230A (Sigma-Aldrich #S8688); TLR4 inhibitor: Viper peptide (Novus#NBP2-226244); CaMKII inhibitor: KN-93 Phosphate (Selleckchem #S7423); DBH inhibitor: Nepicastat (MCE MedChem Express#HY-13289); PKA Inhibitor: 14-22 Amide (Calbiochem®#476485).

### Primary neonatal rat ventricular myocyte (NRVM) and ICA cell isolation

Co-cultured ICA cell-NRVM (NRVM ^ICA+^), NRVM without ICA cell (NRVM ^ICA-^) and ICA cells were isolated and purified using the previously published methods.^24^ Briefly, the neonatal Sprague-Dawley rats (1-3 days) were deeply anesthetized by inhalation of CO_2_, and then the hearts were excised and transferred to pre-cold PBS. (1) NRVM ^ICA+^ isolation procedure: the heart tissue were digested using 0.125% trypsin without EDTA (pH 7.30-7.40) to harvest NRVM ^ICA+^, and continuously cultured in the complete DMEM with 10% FBS, 0.1mM HEPES and 100U/ml Penicillin-Streptomycin for experiments (Supplementary Fig. S1A). (2) NRVM ^ICA-^ purification using superparamagnetic iron oxide particles (SIOP) (BioMag#BM547): NRVM ^ICA+^ suspensions were followed by centrifuging at 800 rpm, 4°C for 7 min. The pellets were resuspended in the pre-cold SIOP solution (40uL SIOP: 4ml PBS), and then used a magnetic separator (Life technologies™) to isolate NRVM ^ICA-^, which contained no ICA cells. (3) ICA cell isolation: the particles in the tube contained SIOP binding ICA cells taken from last step were washed with ice-cold PBS and suspended with complete DMEM. The ICA cells washed down were collected and seeded in plates for further use (Supplementary Fig. S2).

### Langendorff perfusion

Myocardial functions of the adult rat hearts were measured using a Langendorff perfusion system as we described previously.^26^ Briefly, the Sprague-Dawley rats (8-10 weeks) were heparinized (i.p. injection heparin, 2000 U) for 15 min, and then deeply anesthetized with isoflurane inhalation (3% isoflurane in 100% oxygen at a flow rate of 1□L/min) using a face mask. The heart was isolated and the aorta was retrograde set up to a Langendorff perfusion apparatus (Radnoti Langendorff system#120102EZ) with a recirculating mode (volume of 50 mL) to perfuse at 10mL/min with Krebs-Henseleit buffer (bubbled with 95% O2 and 5% CO_2_ gas mixture and maintained at 37 °C). Rat hearts were divided into groups of K-H buffer (control, n=7) and LPS (Sigma-Aldrich, #L2880, Escherichia coli, 055:B5, 1.5 µg/mL) (n=7). The hearts were then performed with a 140 min perfusion of K-H buffer and LPS (1.5 µg/mL) using the recirculating mode, respectively. Collected perfusion fluid at dynamic time points, and harvested tissues at the end of perfusion. In separate experiments, adult rat hearts were arranged into groups of K-H buffer as control (n=7), LPS (n=7) and LPS+Nepicastat (n=5), after a 30-min equilibration period, mixture of LPS (1.5 µg/mL) or/and Nepicastat (a selective DBH inhibitor,^58^ 15 µg/mL) were perfused for 2 h. The physiological parameters of hearts were recorded, and the perfusate and left ventricular tissues were harvested for TNF-α and NE concentration determination as well as immunofluorescence staining (Supplementary Fig. S1F).

### 10X genomics single-cell RNA-sequencing analysis

The strategy for single-cell RNA sequencing analysis of cardiac cells is shown in Fig. 5b. The neonatal Sprague-Dawley rats (1-3 days) were deeply anesthetized with CO_2_ and the hearts were excised and washed in cold PBS to remove the blood. Heart tissues were minced into approximately 1mm^3^ masses and digested using 0.125% trypsin. (1) Cells were prepared using modified Percoll gradient procedure described previously to enrich ICA cells,^59^ of which details are described in Supplementary material. Immuno-staining of the cells was performed to examine the existence of ICA cells prior to single-cell RNA sequencing (Supplementary Fig. S5A). (2) 10X genomics single-cell RNA sequencing: the cells were collected after treatment and analyzed to determine the viability of more than 90% by a cell count system. Individual samples were then loaded on 10X Genomics Chromium System. Cells counted in the system were around 1.1∼1.2*10^4^ per group (Supplementary Fig. S5B). The libraries were prepared following 10X Genomics protocols and sequenced under standard procedure, followed by cell lysis and barcoded reverse transcription of RNA. The library construction and sequencing as well as computational analysis of data were performed at the Saliai Stem cell science company, Guangzhou. Single-cell gene expression was visualized in a two-dimensional projection with *t*-distributed stochastic neighbor embedding (t-SNE) map, where each cell was grouped into one of the 10 clusters (distinguished by their colours), and non-linear dimensional reduction was used (Fig. 5d).^60^ Analyses of batch effect correction and Differentially expressed genes (DEGs) were performed using the Seurat (version: 2.3.4) function RunCCA and FindClusters; Resolution for granularity: 0.5; Differntial expression test: Wilcoxon rank sum test; avg_logFC= log(mean(group1) / mean(group2)); adjusted p_value: Bonferroni Correction; min.pct ≥10%; avg_logFC >= 0.1.

### Immunofluorescence staining

The immunofluorescence staining (IF) was performed according to our previous publication with minor modification.^22^ Briefly, cells were fixed with 4% paraformaldehyde for 15 min. Then, the cells or frozen tissue slides were washed with cold PBS and permeabilized with 0.25% Triton X-100 for 10 min. Subsequently, specimens were blocked at room temperature (RT) for 1 h, and then incubated with primary antibodies at 4°C overnight. On the second day, the specimens were washed with PBS and then incubated in dark at RT with secondary antibodies for 1 h, and then washed twice with cold PBS, followed counterstained with DAPI solution in dark at RT for 10 min. Then the specimens were observed using a laser-scanning confocal microscopy (Leica TCS SP8 X). Details of buffer preparation and antibody dilution are listed in Supplementary Table S2.

### ELISA, Western blotting and Quantitative RT-PCR assay

Norepinephrine (NE) and TNF-α concentration in heart tissues, perfusate and cell supernatants were determined using the NE research enzyme-linked immunosorbent assay (ELISA) kit (ALPCO#17-NORHU-E01-RES) and the TNF-α Quantikine ELISA kit (R&D System#RTA00), respectively. The western blotting assays of proteins were performed using standard protocol.^22^ Antibodies for western blotting assay are available in Supplementary methods. Quantitative RT-PCR assay of relative mRNA expression was analyzed using standard real-time PCR protocol according to TAKARA qPCR kits. In brief, total RNA was reverse transcribed using a PrimeScript™ RT Reagent Kit (TAKARA#RR047A). Real-time PCR were performed with the SYBR Premix Ex Taq II (TAKARA#RR820A) in a LightCycler480 real-time PCR system. Primer sequences for genes are shown in Supplementary Table S3.

### Plasmids, molecular cloning and reporter analysis

The pGL3-Basic Luciferase Reporter vector (Promega), pcDNA3.1 plasmid, plasmids encoding human TRIF and MyD88, RLTK-Luci and AP-1-Luci reporters were a gift from Fuping You (Peking University). For reporter assays, a recombinant pGL3-Rat-TH-Luci reporter was designed and cloned (Supplementary Fig. S3B-D), and then verified by sequencing the relevant region (Supplementary Table S4, Fig. S3E, F). The EGFP plasmid was used as transfection efficiency control (Supplementary Fig. S3G). Hek293/hTLR4-HA cells seeded in 24-well plates were transfected with 50 ng of the luciferase reporter together with a total of 300 ng of various expression plasmids or empty control plasmids. Quantification of dual luciferase activity was performed using a GALEN dual-luciferase assay kit (GALEN#GN201).

### RNA interference

RNA interference was designed to disrupt the AP-1 binding in rat TH promoter region (Supplementary Fig. S4A-C). c-Jun siRNA, c-Fos siRNA (Sequences refer to Supplementary Table S5), and control siRNA transfection was processed using Lipofectamine^®^3000 Reagent (Invitrogen™#L3000001) and Opti-MEM (Gibco#31985-070) under standard protocol.

### Isolation of primary rat peritoneal macrophages

Rat peritoneal macrophages were isolated using published method.^61^ Adult rats (250-300 g) were euthanized with CO_2_ plus cervical dislocation, and then the abdomen was soaked with 70% alcohol followed by making a small incision along the midline. 10 ml of DMEM were injected into each rat abdomen and gently massaged. About ∼8 ml fluid was aspirated from peritoneum per rat. The peritoneal cells were collected by centrifuging for 10 min, 400×g at 4°C.

### Statistical analysis

All data are reported as mean ± standard error of mean (SEM), where *n* is the number of animals or independent experiments. Statistical analyses were conducted using SPSS 13.0 (SPSS Inc., Chicago, IL, USA). The statistical significance of the difference between the two groups was measured by a two-tailed independent Student’s *t-test* or ANOVA with Bonferroni post hoc analysis, as appropriate. Statistical significance was accepted at *P* < 0.05.

## Acknowledgments

We thank Fuping You in Peking University Health Science Center for the plasmids. We thank Bo Yu and Quan Yu in the Medical Experimental Center of School of Medicine at Jinan University for expert technical assistance in using confocal microscopy. This work was supported by grants from the National Natural Science Foundation of China (81871542).

## Author contributions

Duomeng Yang and Huadong Wang conceived the study and designed the experiments. Duomeng Yang, Xiaomeng Dai and Yun Xing performed the majority of the experiments and analyzed the data; Xiangxu Tang, Guang Yang, Hongmei Li, Xiuxiu Lv and Xiaohui Yu contributed specific experiments and data analysis. Penghua Wang, Andrew G. Harrison, Duomeng Yang and Huadong Wang wrote the paper.

## Declaration of interests

The authors declare no competing interests.

## Notes

### Competing Interest Statement

The authors have declared no competing interest.

