## Supplemental Fig S1-S6 and Table S1-S5 for "Intrinsic cardiac adrenergic cells contribute to septic cardiomyopathy"

**Title page**

Running title: ICA cell facilitates septic cardiomyopathy

***Corresponding author:** Prof. Huadong Wang, MD, PhD,, Department of Pathophysiology, School of Medicine, Jinan University, Guangzhou 510632, Guangdong, China. Tel: 86-20-85220241; Fax: 86-20-85221343. ORCID: 0000-0002-2197-3624

### 1. Supplementary figures/Tables

#### 1.1 Supplementary figures

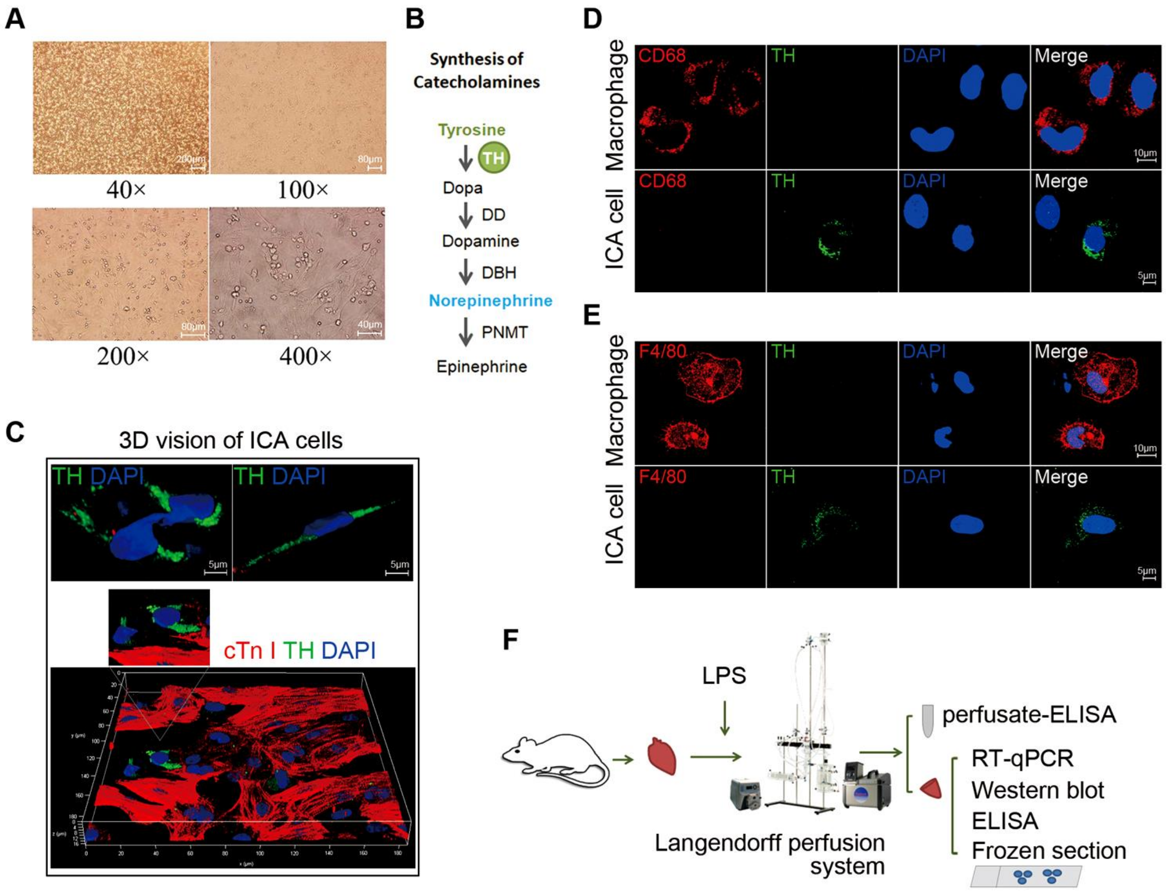


**Fig. S1 ICA cells in primary cultured neonatal rat cardiomyocytes and Langendorff perfused adult rat hearts.** (*A*) Primary co-cultured ICA cell-NRVM (NRVM $^{ICA+}$) isolated using traditional enzymatic method. Cells are from *n*=6 neonatal rats. (*B*) Catecholamine biosynthetic pathway. TH, tyrosine hydroxylase; DD, DOPA decarboxylase; DBH, dopamine β-hydroxylase; PNMT, phenylethanolamine N-methyltransferase. (*C*) 3D vision of ICA cells. Cardiac troponin I (cTn I): cardiomyocytes, red; TH: ICA cells, green; DAPI: nuclei, blue. Cells are from *n*=6 neonatal rats. (*D* and *E*) ICA cells and macrophages, CD68 and F4/80: macrophages, red; TH: ICA cells, green. ICA cells are from *n*=6 neonatal rats. Peritoneal macrophages are from *n*=2 adults rats. (*F*) Langendorff perfusion system for isolated adult rat hearts.

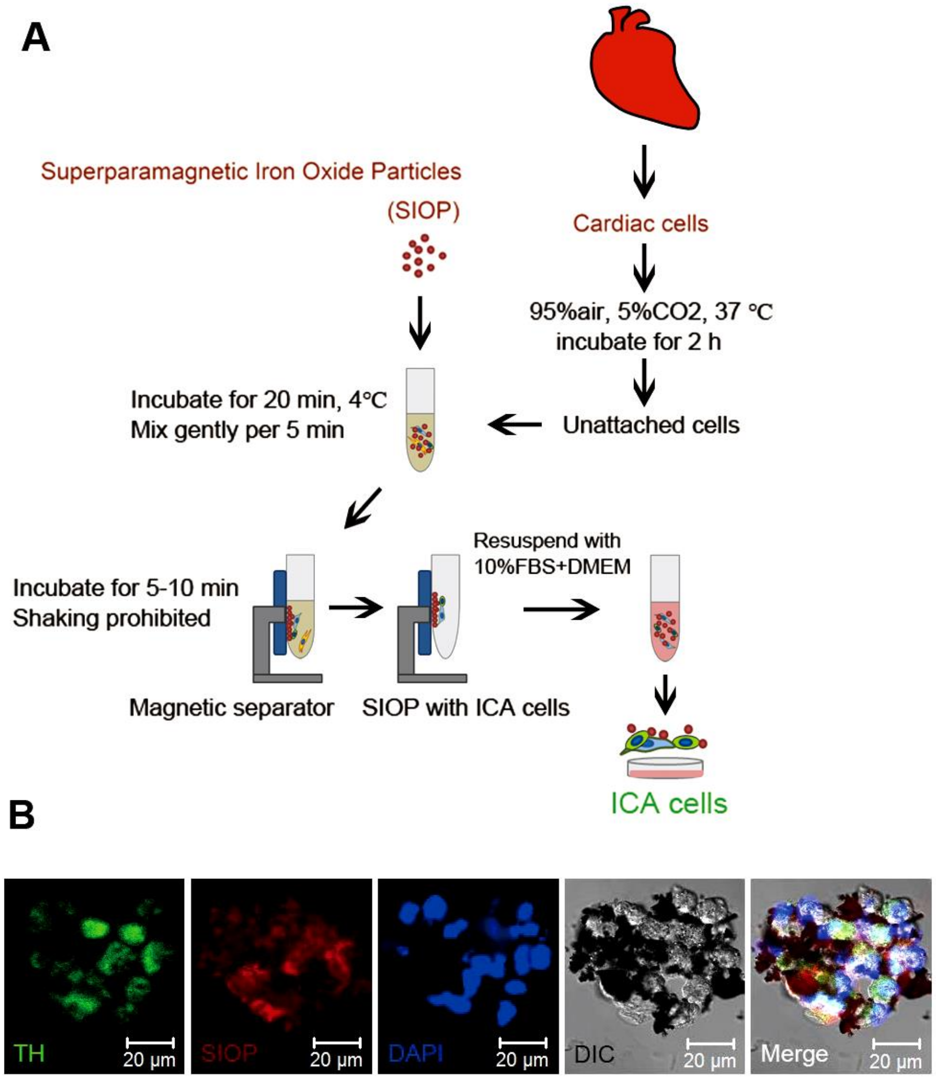


**Fig. S2 Procedure of ICA cell isolation using superparamagnetic iron oxide particles (SIOP).** (*A*) Schematic procedure of ICA cell isolation using SIOP. (*B*) Immuno-staining of ICA cells binding to SIOP. Cells are from *n*=12 neonatal rats.

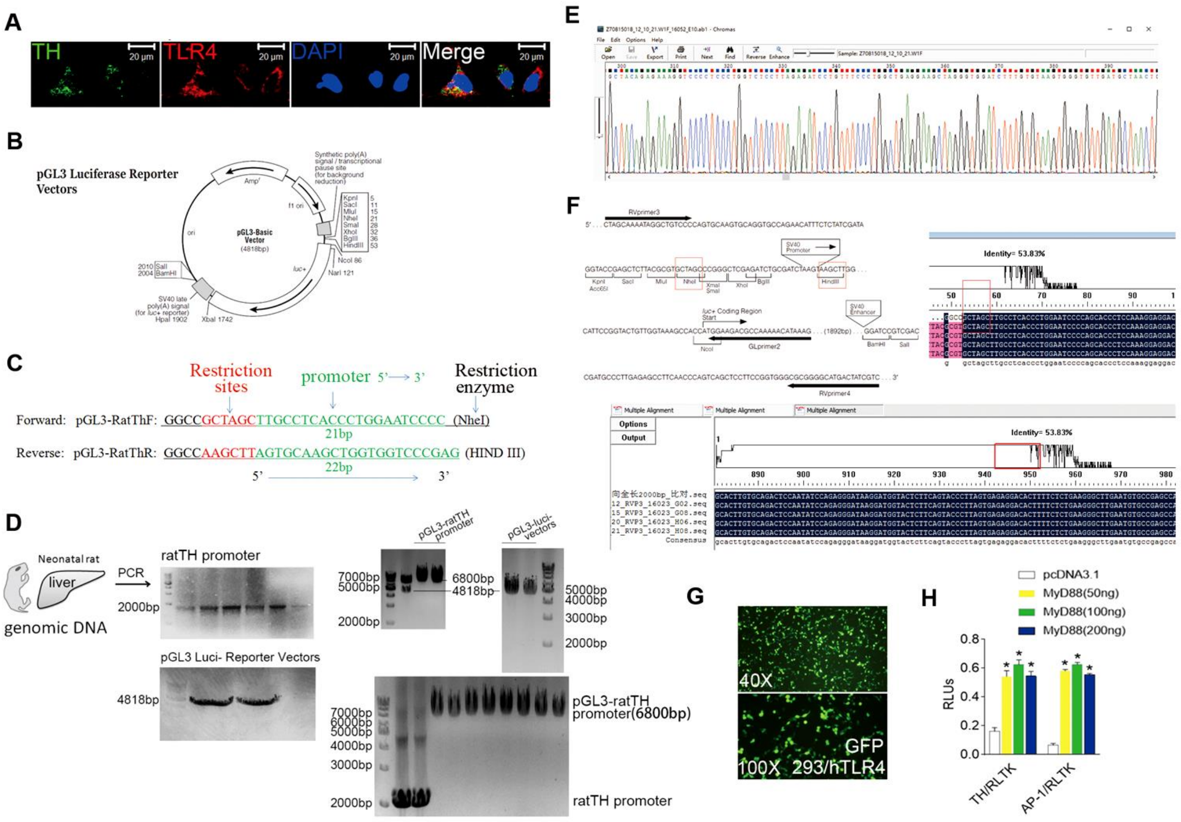

**Fig. S3 TLR4 expression on ICA cells and recombination of pGL3-luc- rat TH promoter.** (*A*) Immuno-staining of TLR4 expressed on ICA cells purified using SIOP. TLR4: toll-like receptor 4, red; TH: ICA cells, green; DAPI: nuclei, blue. ICA cells are from *n*=12 neonatal rats. (*B*) pGL3-Basic luciferase reporter Vector. (*C*) Design of rat TH promoter primers. (*D*) Cloning of pGL3-luc- rat TH promoter (6800bp). The liver tissue was from a neonatal rat. (*E* and *F*) Sequence of the pGL3-luc- rat TH promoter. (*G*) EGFP expression as a positive control 24 h after plasmid transfection. *n*=3 independent experiments, cell density=3*$10^5$ cells/mL. (*H*) TH-luci expression relative to RLTK stimulated by different dose of MyD88 stimulation. Data are from *n*=3 independent experiments, cell density=3*$10^5$ cells/mL. Data are presented as mean ± S.E.M. and analyzed using one-way ANOVA with Bonferroni post hoc test or two-tailed independent Student's t-test as appropriate, **P*<0.05, ***P*<0.01. Experiments performed in *n*=3 independent cell isolations are biological repilicats.

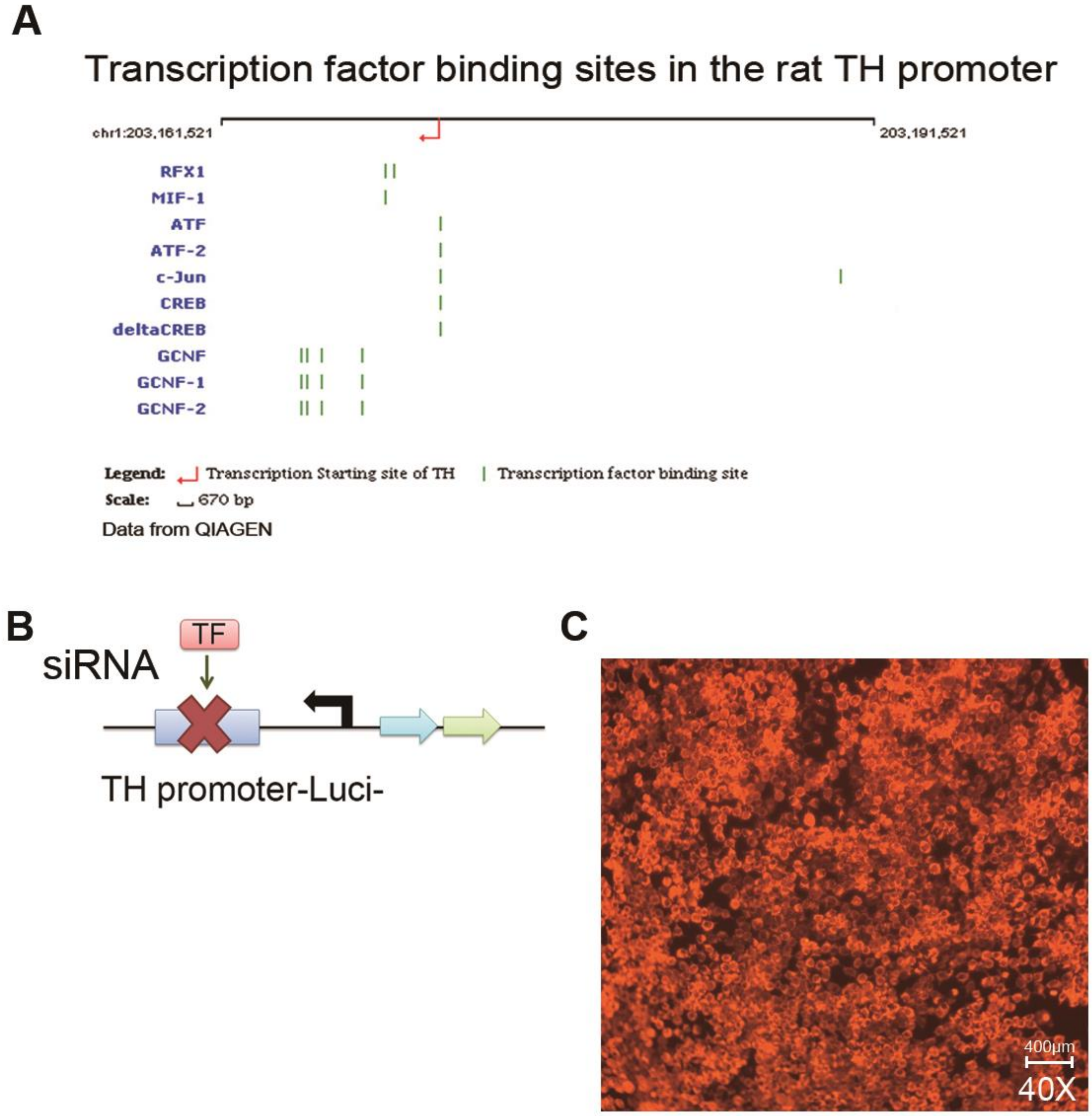


**Fig. S4 Strategy for determining transcription factors mediating TLR4 activation of TH promoter.** (*A*) Transcription factor binding sites in the rat TH promoter. (*B*) siRNA strategy designed to disrupt activation of TH promoter. (*C*) Fluorescence of Cy3-control siRNA 24 h after transfection into 293/hTLR4 cells. *n*=3 independent experiments, cell density=3*$10^5$ cells/mL.

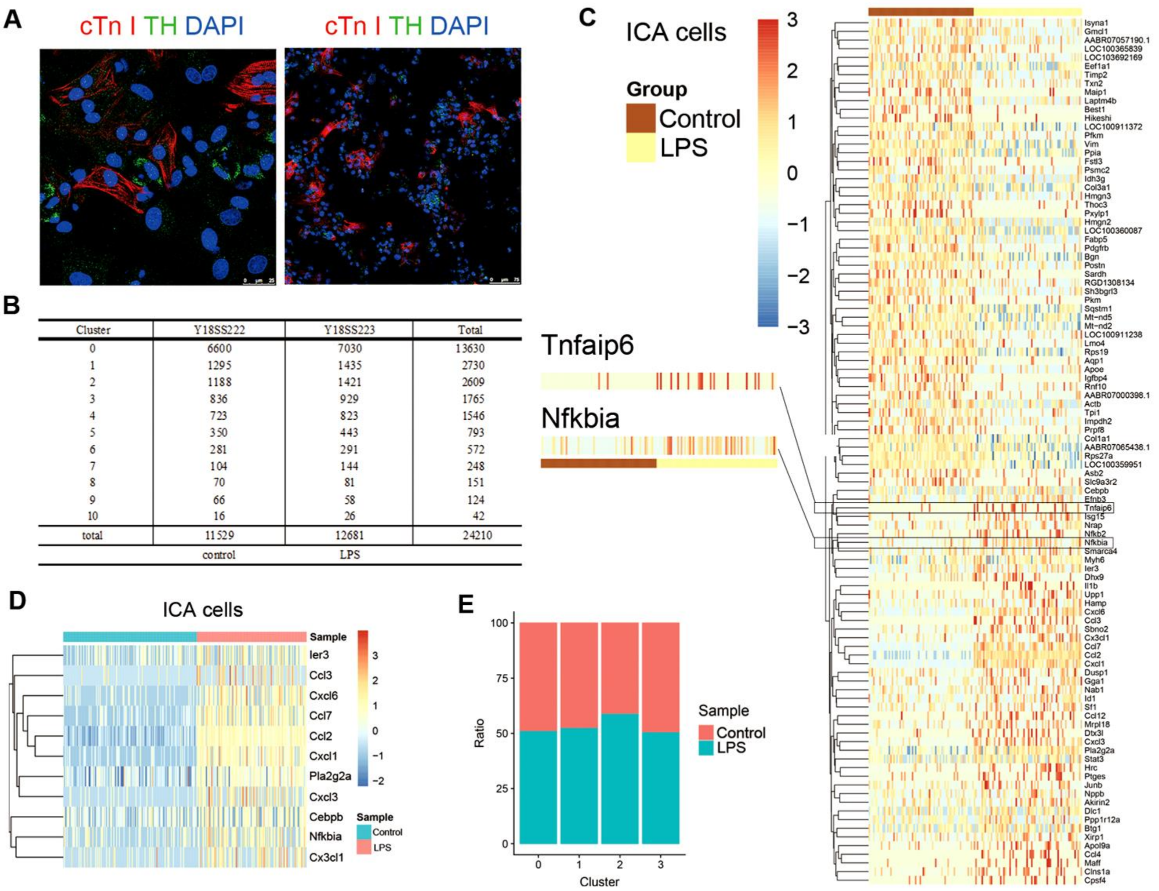


| Cluster | Y18SS222 | Y18SS223 | Total |
|---|---|---|---|
| 0 | 6600 | 7030 | 13630 |
| 1 | 1295 | 1435 | 2730 |
| 2 | 1188 | 1421 | 2609 |
| 3 | 836 | 929 | 1765 |
| 4 | 723 | 823 | 1546 |
| 5 | 350 | 443 | 793 |
| 6 | 281 | 291 | 572 |
| 7 | 104 | 144 | 248 |
| 8 | 70 | 81 | 151 |
| 9 | 66 | 58 | 124 |
| 10 | 16 | 26 | 42 |
| total | 11529 | 12681 | 24210 |
| | control | LPS | |

**Fig. S5 10X genomics single-cell RNA sequencing.** (*A*) Immuno-staining of cardiac cell samples prior to single-cell sequencing processing. Cardiac troponin I (cTn I): cardiomyocytes, red; TH: ICA cells, green; DAPI: nuclei, blue. (*B*) Number of Cells counted for different cluster Single-cell transcriptomes; (*C*) Heatmap of Top 100 differentially expressed genes (DEGs) in ICA cells between control and LPS groups. (*D*) Top variable DEGs in the ICA cells between control and LPS groups, control: saline. (*E*) Ratio of cells in different subclusters of ICA cells and cardiomyoyte cluster. Analysis of batch effect correction and DEGs were performed using the Seurat (version: 2.3.4) function RunCCA and FindClusters; Resolution for granularity: 0.5; Differntial expression test: Wilcoxon rank sum test; avg_logFC= log(mean(group1)/mean(group2)); adjusted p value: Bonferroni Correction; min.pct ≥10%; avg_logFC >= 0.1. Cells for single cell RNA-seq are from *n*=24 neonatal rats.

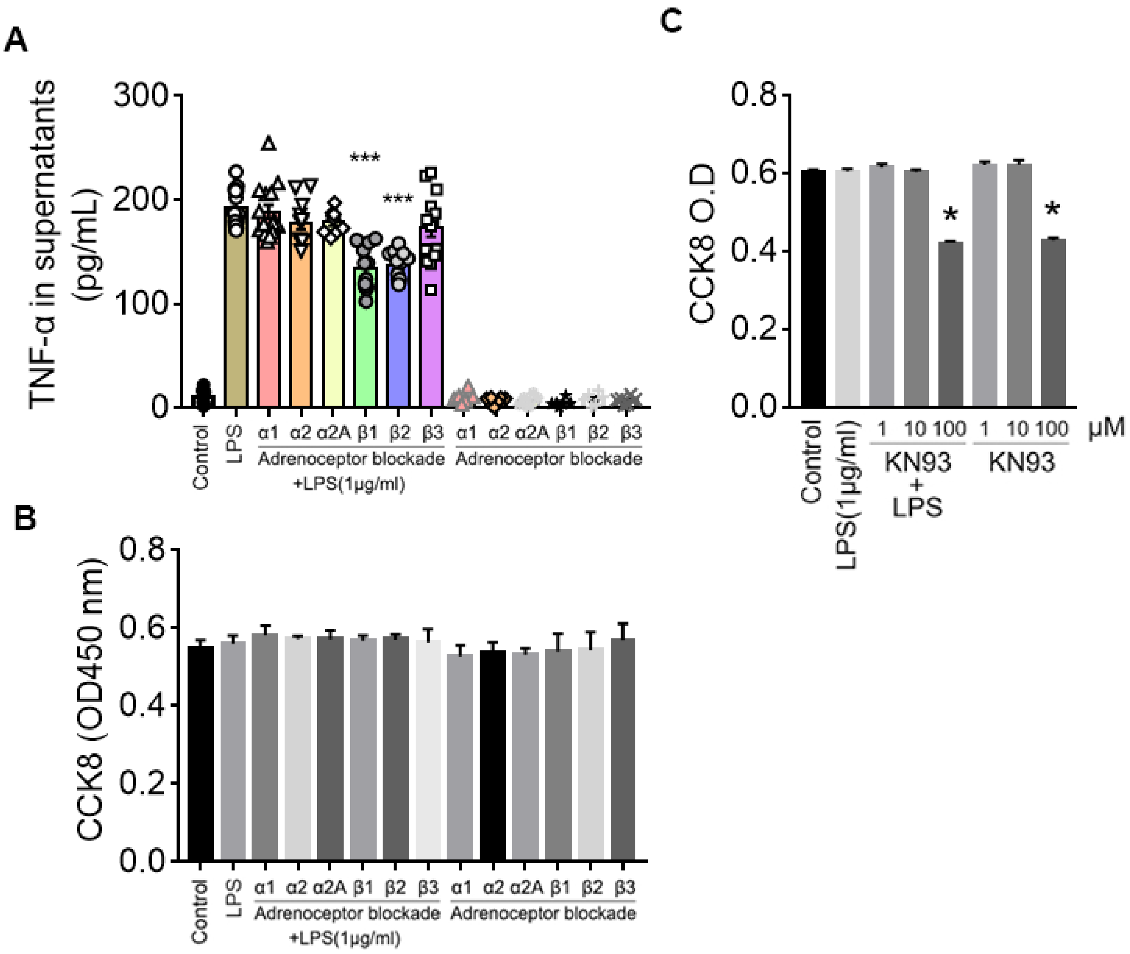


**Fig. S6 Effects of different adrenergic receptor blockade on LPS-induced TNF-α production.** (*A*) Effect of different adrenoceptor blockers on NRVM $^{ICA\ cell+}$ TNF-α production induced by LPS. Data were performed in *n*=6 independent experiments with biological triplicate each time. (*B*) CCK8 assay of NRVM $^{ICA\ cell+}$ treated with different adrenoceptor blockers for 30 min prior to LPS administration. Data were performed in *n*=3 independent experiments with 5 biological replicats each time. (*C*) CCK8 assay of NRVM $^{ICA\ cell+}$ treated by KN93 for 30 min prior to LPS administration. Data were from *n*=3 independent experiments. Data are presented as mean ± S.E.M. and analyzed using one-way ANOVA with Bonferroni post hoc test, **P*<0.05, ****P*<0.001.

### 1.2 Supplementary Tables

Table S1

Preparation of discontinuous Percoll gradients solution

| Percoll solution | 12.5×Density buffer (mL) | Sterile water (mL) | Percoll (mL) | Final volume (mL) |
| --- | --- | --- | --- | --- |
| Low density (1.060 g/mL) | 1.04 | 6.64 | 5.32 | 13 mL |
| High density (1.086 g/mL) | 1.04 | 4.02 | 7.94 | 13 mL |

Table S2

Preparation of buffer solution and antibody dilution for IF

| | |
|---|---|
| Blocking buffer | 1% BSA, 10% donkey serum, 0.3M Glycine and 0.25% Triton X-100 in PBS |
| Antibody and DAPI dilution buffer | 1% BSA, 0.3M Glycine and 0.25% Triton X-100 in PBS |
| Antibody dilution | Anti-Tyrosine Hydroxylase antibody (MerckMillipore#AB152, dilution 1:200);<br>Anti-Vimentin antibody (Abcam#ab24525, dilution 1:200);<br>Anti-TLR4 antibody (Abcam#ab22048, dilution 1:200)<br>Anti-Cardiac Troponin I antibody (Abcam#ab10231, dilution 1:200);<br>Anti-CD68 antibody (Abcam#ab201340, dilution 1:200);<br>Anti-F4/80 antibody (Abcam#ab186073, dilution 1:200);<br>Donkey anti-rabbit IgG (H+L) highly cross-adsorbed secondary antibody Alexa Fluor®488 (Invitrogen™#A-21206, dilution 1:200);<br>Goat anti-Chicken IgY/Alexa Fluor 647 (Bioss#bs-0310G-AF647, dilution 1:200);<br>Donkey anti-Mouse IgG (H+L) highly cross-adsorbed Alexa Fluor®555 (Invitrogen™ #A-31570, dilution 1:200);<br>Donkey anti-Rabbit IgG H&L (Alexa Fluor® 594) preadsorbed (Abcam#ab150064, dilution 1:200) Goat anti-Chicken IgY H&L (Alexa Fluor®488) preadsorbed (Abcam#ab150173, dilution 1:200); |

Table S3

Primer sequences for RT-qPCR assays

| mRNA | Primer | Sequences (5'--3') |
|---|---|---|
| TH | Forward primer | GTCTCAGAGCAGGATGCCAAG |
| | Reverse primer | AGAACAGCATTCCCATCCCT |
| DBH | Forward primer | CTGGTGTACACGCCCTTGAT |
| | Reverse primer | AGGCAAAGATGCGGATTCCA |
| GAPDH | Forward primer | GGCACAGTCAAGGCTGAGAATG |
| | Reverse primer | ATGGTGGTGAAGACGCCAGTA |

Table S4

Sequence of pGL3-Rat-TH promoter luciferase reporter plasmid

| Sequence (promoter regions -2000-+1, 5’-3’) |
|---|
| TTCTCTATCGATAGGTACCGAGCTCTTACGCGTGCTAGCTTGCCTCACCCTGGAATC |
| CCCAGCACCCTCCAAAGGAGGACCCTGGGAGTGGGCATAGACGCCCTTCAGGTG |
| TGGGCAACAGCCCCCAGTCCTCAGGATGAAAGGCTAAGGTGCAGCCAGCTCTGC |
| CTTCACGGTGGGAATGTCTCTATGTGAGCCCTTTCTGGGCTGTGAAGAACGCTCTG |
| AGAAGGGTCCTGGGACCCTGGATAGGCCAGAGCTGTGCTGGGCATGTAGAGACA |
| GGAGTGGGCTAAAGCAGCAAAGGCACTGACCAAGGAAGAGTTCAGAGAGGAGC |
| GTGGAATATGGGGAGGGGTTCATAGTAAGAGAGAGCAGGCAGTGGAGAGTAAATA |
| GTCACTGAGCCGGGGTTTATGGGGTTTGTAGGAGCTTACTCAGAGAAAGTAGATG |
| AGAGATGCCATGCCAGTCTGAGTATCACAGAGCCCCAGGCTCTCCTGGGAACGGA |
| ACTGTGAGGGCCAGAAGGTCAGCAAGGGAGGTTAGGGAGAGTTCCTTTTGTACT |
| GACTCAGCATTTATCCTGCTCCCAGGGGGCAATGGGGGCCAGTGAGGGATGCAGA |
| GCAAGGCAGTGATGTGGCAGGCAGTTCCTGTTGTGAAAGAGCTGGGAAGGGAGC |
| GGGCTGGGCCTGGTACGTACAGCAGGCCATTTCTGAGGGTCCGAGTGCTGTCTAG |
| GAGGTGCAGTGAGACTTCAGTGATCAGCCAGAACAGAAGCTAAGCGGGGTGGGG |
| ACTGCGAGTTCAGGCTTCTGGGTCTTGCAAATATCCAGAATGCTAAATCCTCAGAA |
| CCCCAGGGTGGCCATTTTCAGAGTGGGTTTTGTCCTTTGGGCACTTGTGCAGACTC |
| CAATATCCAGAGGGATAAGGATGGTACTCTTCAGTACCCTTAGTGAGAGGACACTT |
| TTCTCTGAAGGGCTTGAATGTGCCGAGCCATTACCTGAAGGAAGGAAATGACTCC |
| AGGGACATAAGATGGGCCCAGCACAACTCACCTGCTACAGAGAAAGGTCCCCTCC |

CTGGTCTCCTTAGAGATCCTGTTTCCCTGGCTGAGGAAGCTAGGGTGGATCTTTGT

GTAAGTGGGTGTTGATGCTAACTGGAAAACAAAAGGTCACTTACCGTTAGACCTC

GGGGTACCATGGAAGAGATGATCACTGAGTGTGCCCTTACATGGGGACCAGCTGA

GAATGGGGCTACCACTAGCTCGAGACCATGATACAGGGAATAAGTGTGCATTTGG

GGGTAGGGAGTGGCTCAGAATACTCTTAACCAAAGCAGAGGTTTGCTCCCACAGG

AAGGTGAGGTCAGAAGGCCTTAGGGAGCTGCCAGGGGCTAGGGTTGGCACCATC

TCCCAGGCTGTGTCTTTAAGGAGATGATAATCAGAGGGATAGAACCTTGCAAAAG

TGGGCCAGTCTTGGGAATACTATAGAGGAATAGCCTTCTGGAACATTCTGTGTCTC

ATAGGACCTGCCTGGGGATCCAGCCCCAGTGCCAGCACATATACCGACTGGGGCA

GTGAATAGATAGTACACTTTGTTACATGGGCTGGGGGGAACATGGCCCATGTCCTG

GAGGGGACTTTATGACAGACATCCAAAAATCCAGTGAGAGGGCTTCTAGATTTGT

CTCCAAAGGTTATAGTTCTAACATGAGCCCTTAGGAAATCCAGCATAGTTCTCCCT

GTGTGCCCTGGTTTGGTTAGAGAGCTCTAGCGGTCTCCTGTCCCACAGAATACCAG

CCAGCCCCTGCCCTACGTCGTGCCTCGGGCTGAGGGTGATTCAGAGGCAGGTGCC

TGTGACAGTGGATGCAATTAGATCTAATGGGACGGAGGCCTTTCTCGTCGCCCTCG

CTCCATGCCCACCCCCGCCTCCCTCAGGCACAGCAGGCGTGGAGAGGATGCGCAG

GAGGTAGGAGGTGGGGGACCCAGAGGGGCTTTGACGTCAGCCTGGCCTTTAAAG

AGGGCGCCTGCCTGGCGAGGGCTGTGGAGACAGAACTCGGGACCACCAGCTTGC

ACTAAGCTTGGCATCC

Table S5

| | |
|---|---|
| c-Jun siRNA | stB0003638A, sequence: CCAACATGCTCAGGGAACA; |
| | stB0003638B, sequence: GGGTGCCAACTCATGCTAA; |
| | stB0003638C, sequence: TGGAGCGCCTGATAATCCA; |
| c-Fos siRNA | stB0003606A, sequence: GGGATAGCCTCTCTTACTA; |
| | stB0003606B, sequence: CCTGCAAGATCCCTGATGA; |
| | stB0003606C, sequence: GACCTATCTGGGTCCTTCT |

### 2. Supplementary Methods

#### 2.1 Animals and cells

All experiments with animals were conducted in compliance with the Guide for the Care and Use of Laboratory Animals published by the US National Institutes of Health and approved by the Animal Care and Use Committee at Jinan University (LAECJU20180225016). For Langendorff perfusion, the rats were deeply anaesthetized with isoflurane (3% isoflurane in 100% oxygen at a flow rate of 1 L/min) using a face mask. For primary neonatal cardiomyocyte isolation and peritoneal macrophage isolation, the rats were euthanized with carbon dioxide ($CO_2$) plus cervical dislocation.

The neonatal (1-3 days old) and adult (6-8 weeks) Sprague-Dawley rats were obtained from the laboratory animal center of Southern Medical University (Guangzhou, China). The TLR4-deficient mice ($Tlr4^{Lps\text{-}del}$, strain: C57BL/10ScNJNju) were purchased from Nanjing Biomedical Research Institute of Nanjing University. Neonatal rat ventricular myocyte (NRVM). ICA cells and peritoneal macrophages were primarily isolated from rats or mice. 293/hTLR4-HA cells (InvivoGen#293-htlr4ha) were a gift from Guang Yang (Jinan University) and designed for studying the stimulation of hTLR4. All of the cell culture plates and dishes were purchased from BIOFIL® (Guangzhou, China).

#### 2.2 Inhibitors

TLR4 inhibitor: VIPER Peptide (Novus#NBP2-226244); $\alpha_1$-AR antagonist: Prazosin (Merck Sigma-Aldrich#7791); $\alpha_2$-AR antagonist: Yohimbine (Merck Sigma-Aldrich #Y3125); $\alpha_{2A}$-AR antagonist: BRL 44408 (Merck Sigma-Aldrich #B4559); $\beta_1$-AR antagonist: CGP20712A (Merck Sigma-Aldrich #C231); $\beta_2$-AR antagonist: ICI-118 551 (Merck

Sigma-Aldrich #I127); $\beta_3$-AR antagonist: SR59230A (Merck Sigma-Aldrich #S8688); CaMKII inhibitor: KN-93 Phosphate (Selleckchem #S7423); PKA Inhibitor 14-22 Amide (MerckMillipore Calbiochem®#476485); Dopamine-β-hydroxylase (DBH) inhibitor: Nepicastat (MCE MedChem Express#HY-13289)

**2.3 Primary neonatal rat ventricular myocyte (NRVM) and ICA cell isolation and culturing**

Co-cultured ICA cell-NRVM (NRVM $^{ICA+}$), NRVM without ICA cells (NRVM $^{ICA-}$) and ICA cells were isolated and purified using the previously published method.[2] The neonatal Sprague-Dawley rats (1-3 days) were deeply anesthetized with carbon dioxide ($CO_2$) and sacrificed by cervical dislocation, and then the hearts were excised and transferred to pre-cold PBS (HyClone™#SH30256.01). The whole procedure was performed in three steps:

**1) NRVM $^{ICA+}$ isolation procedure**

The excised hearts from neonatal rats were put into cold PBS and washed 3 times to remove the blood. Cut one third from apex of the heart, these ventricular tissues were put into a new dish and washed again. Then the tissues were minced into approximately 1mm$^3$ masses which were subsequently put in the 0.125% trypsin buffer without EDTA and phenol red (Gibco®#15090046, pH7.30-7.40). Suspended the tissue pieces and performed digestion in a 37°C water bath with magnetic stirring for 7 minutes at low speed (<100 rpm). The supernatant of this first digestion was discarded, and then re-add 0.125% trypsin buffer to repeat this digestion process. Collected the digestion supernatant, added equal volume DMEM (HyClone™#SH30243.01) with 10% FBS (BOVOGEN#SFBS-AU) and mixed softly to stop enzymatic digestion. Repeated this digestion step 5-6 times until few residual light-color tissues and collected totally 5-6 tubes digestion supernatants contained cardiac cells. The supernatants were centrifuged at 800 rpm for 7 minutes at 4°C. The cell pellets were resuspended softly with 5ml DMEM and washed one more time. Finally, softly suspended cell pellets with 6ml complete DMEM (DMEM with 10% FBS, 0.1mM HEPES (Sigma-Aldrich#V900477) and 100U/ml Penicillin-Streptomycin (HyClone™#SV30010)) and then filtered through a 70μm cell strainer into a 25cm$^2$ cell culture flask (for less than 20 neonatal rats). The cells were then incubated in 95% air and 5% $CO_2$ at 37°C for 2h to remove most non-cardiomyocytes. After this 2-hour differential attachment, collected supernatants containing unattached cardiac cells in a new 50ml centrifuge tube and mixed the cells suspension softly. These cardiac cells contained certain amount of ICA cells during culture, therefore were considered as co-cultured ICA cell-NRVM (NRVM $^{ICA+}$). NRVM $^{ICA+}$ were plated and cultured for an appropriate length of time, and then used for experiments (*Figure S1A*).

**2) NRVM $^{ICA-}$ purification using Superparamagnetic iron oxide particles (SIOP)**

NRVM $^{ICA-}$ was purified from fresh NRVM $^{ICA+}$ by using SIOP (BioMag#BM547). NRVM $^{ICA+}$ suspensions were pipette into a 15ml centrifuge tube followed by centrifuging at 800 rpm for 7 min at 4°C. Discarded supernatants, the cell pellets were resuspended in pre-cold label-free SIOP solution which was diluted with the ratio of 40uL SIOP: 4ml PBS, and then incubated at 4°C for 20 min with gentle mixing every five minutes. After incubating, the tube containing cells and SIOP mixture was set up into a magnetic separator (Life technologies™) standing for 5-10minutes until the SIOP were totally stuck to the wall of tube and the supernatant was clear. The supernatant was carefully transferred into another 15ml tube by pipette and centrifuged at 800 rpm for 7 minutes at 4°C. Discarded the supernatant, the cell pellets were resuspended in certain volume of culture medium. These cells were NRVM $^{ICA-}$ of which purity was enriched up to more than 93%, and importantly contained no ICA cells during culture.[2] NRVM $^{ICA-}$ were then plated and cultured for an appropriate length of time for further experiments

**3) ICA cells isolation**

A schematic procedure of the methodology for ICA cell isolation is shown in *Figure S2*. The particles in the tube contained SIOP binding ICA cells taken from step (2) were washed with ice-cold PBS and suspended with complete DMEM. This ICA cell suspension was collected in certain volume and seeded in culture plates for further use.

**2.4 Langendorff perfusion**

Myocardial functions of the hearts isolated from adult rats were measured using a Langendorff perfusion system as we previously described.[3] Briefly, the Sprague-Dawley rats (8-10 weeks, 250-300 g) were heparinized (i.p. injection heparin, 2000 U) for 15 min, and then deeply anesthetized with isoflurane inhalation (3% isoflurane in 100% oxygen at a flow rate of 1 L/min). The hearts were isolated, and then the aorta was retrograde set up to a Langendorff perfusion apparatus (Radnoti Langendorff system#120102EZ) to perfuse at 10mL/min with Krebs-Henseleit buffer containing (in mM) 118 NaCl, 4.7 KCl, 25 $NaHCO_3$, 1.2 KH2$PO_4$, 1.2 $MgSO_4$, 2.5 $CaCl_2$ and 11 glucose (bubbled with 95% O2 and 5% $CO_2$ gas mixture and maintained at 37 °C). A balloon was inserted into the LV chamber through the mitral valve with an incision in the left atrium and connected to a pressure transducer for the continuous measurement of left ventricular (LV) pressure. The LV balloon volume was adjusted to approximately 10 mmHg of the LV end-diastolic pressure for stabilization, and the left ventricular developed pressure and the maximum rates of positive and negative changes in the LV pressure (±dP/dt) were calculated using a biological signal acquisition and processing system (Tai Meng#BL-420F).

Adult rat hearts were divided into two groups: K-H buffer as control (n=7) and LPS (Sigma-Aldrich, #L2880, Escherichia coli, 055:B5, 1.5 μg/mL) (n=7). The hearts were set up on the Langendorff system with a recirculating mode with K-H buffer (total volume, 50 mL), and then were performed with a 140 min perfusion of K-H buffer and LPS (1.5 μg/mL), respectively. The perfusion fluid was collected at different time points, and left ventricular tissues were harvested at the end of perfusion for mRNA and protein expression determination. In separate experiments, adult rat hearts were arranged into groups of K-H buffer as control (n=7), LPS (n=7) and LPS+Nepicastat (n=5), and placed on the Langendorff apparatus with perfusing in a recirculating mode with Krebs-Henseleit buffer (total volume, 50 mL). After a 30-min equilibration period, LPS (1.5 μg/mL) or/and Nepicastat (a selective DBH inhibitor,[4] 15 μg/mL, MCE MedChem Express #HY-13289) mixed in the K-H buffer were perfused for 2 h. The above physiological parameters of hearts were recorded, and the perfusate and left ventricular tissues were harvested for TNF-α and NE concentration determination as well as Immunofluorescence staining (*Figure S1F*).

**2.5 Isolation of primary rat peritoneal macrophages**

Rat peritoneal macrophages were isolated using previously methods with minor modification.[5] Adult rats (8-10 weeks, 250-300 g) were euthanized with carbon dioxide ($CO_2$) plus cervical dislocation. The abdomen was soaked with 70% alcohol and then made a small incision along the midline with sterile scissors. 10 ml of DMEM were injected into each rat abdomen and gently massaged. A syringe with needle was used to aspirate fluid from peritoneum. About ~8 ml fluid recovery per rat were expected. The peritoneal cells were collected by centrifuging for 10 min, 400×g at 4°C. Supernatants were discarded and cell pellets were resuspended in cold DMEM by gently pipetting up and down. The cell density was then adjusted in DMEM for further use.

**2.6 Single-cell RNA-sequencing analysis**

The strategy for single-cell RNA sequencing analysis of cardiac cells is shown in *Figure 5B*.

(1) Cell preparation using Percoll®

Cells for Single-cell RNA sequencing were prepared using modified Percoll gradient procedure described by others previously to enrich ICA cells.[6] Percoll density gradient buffer (12.5×) was prepared: NaCl 8.47 g, HEPES 5.96 g, $NaH_2PO_4$ 0.17 g, glucose 1.24 g, KCl 0.5 g, $MgSO_4$ 0.25, dissolved in a final volume of 100 mL of deionized water, pH to 7.4 and filter sterilize. Discontinuous Percoll gradients were prepared as description in *Table S1***.**

The neonatal Sprague-Dawley rats (1-3 days) were deeply anesthetized with carbon dioxide ($CO_2$) and sacrificed by cervical dislocation. The hearts were excised and put into cold PBS, washed 3 times to remove the blood. Cut one third from apex of the heart, these ventricular tissues were minced into approximately 1mm$^3$ masses and digested using 0.125% trypsin. After getting single cardiac cell suspension from digestion, created a two layer density gradient by carefully layering 3 mL of the low-density Percoll solution (1.060 g/mL) on the top of 3 mL of the high-density Percoll solution (1.086 g/mL) to form a discontinuous gradient. The cell suspension (2 mL) is then layered on the top of the 1.060 g/mL Percoll solution. Centrifuged at 1800 ×g for 45 min, at room temperature, the non-NRVMs (including cardiac fibroblasts, immature cardiomyocytes, ICA cells, some cardiomyocytes and other cells) were located at the upper low-density gradients. The Percoll band containing non-NRVMs were harvested using a transfer pipet, and then added medium to make a final volume of 10 mL. Centrifuged (1800 ×g, 10 min, 25 ℃), and 5 mL of medium was added to resuspended the cell pellets. Cells were counted, plated, cultured and treated with LPS or saline (control). Prior to 10X genomics single-cell RNA sequencing, the cells obtained from Percoll procedure were made immuno-staining to make sure the existence of enriched ICA cells (*Figure S5A*).

(2) 10X genomics single-cell RNA sequencing

After treatment, cells were digested by 0.125% trypsin with EDTA and collected in complete DMEM. The cell viability was up to more than 90% determined by a cell count system. Individual samples were loaded on 10X Genomics Chromium System. Cells counted in system were around 1.1~1.2*$10^4$ per group (*Figure S5B*). Libraries were prepared following 10X Genomics protocols and sequenced under standard procedure, followed by cell lysis and barcoded reverse transcription of RNA. The library construction and sequencing as well as computational analysis of data were performed at the Saliai Stem cell science company, Guangzhou. Single-cell gene expression was visualized in a two-dimensional projection with t-SNE, where each cell is grouped into one of the 10 clusters (distinguished by their colours), and non-linear dimensional reduction is used (*Figure 5D*).[7] Analysis of batch effect correction and Differentially expressed genes (DEGs) were performed using the Seurat (version: 2.3.4) function RunCCA and FindClusters; Resolution for granularity: 0.5;

Differntial expression test: Wilcoxon rank sum test; avg_logFC= log(mean(group1) / mean(group2)); adjusted p_value: Bonferroni Correction; min.pct ≥10%; avg_logFC >= 0.1.

**2.7 Immunofluorescence staining**

We performed immunofluorescence staining (IF) according to our previous publication [8] with minor modification. Briefly, cells were cultured as experiment request; the heart tissues were harvested, fixed in 4% paraformaldehyde, infiltrated with Tissue Tek OCT compound (SAKURA#4583) and rapidly frozen to -80°C before sectioning. Cells in confocal dishes were washed twice by cold PBS and fixed with 4% paraformaldehyde for 15 minutes. Then, the cells or frozen tissue slides were washed three times with cold PBS and permeabilized with 0.25% Triton X-100 in PBS for 10 minutes followed by washing three times. Subsequently, cells or frozen tissue slides were blocked at room temperature for 1 hour. After blocking, cells or frozen tissue slides were then incubated with primary antibodies at 4°C overnight. After three-time wash in PBS, the frozen tissue slides or cells were incubated in dark with secondary antibodies for 1 h at room temperature, and then washed twice with cold PBS, followed counterstained with DAPI solution (Dojindo#D523, dilution 1:200) in dark for 10 minutes at room temperature. Then cells or frozen tissue slides were observed by a laser-scanning confocal microscopy (Leica TCS SP8 X, Leica Microsystems). Buffer preparation and antibody dilution details are listed in *Table S2*.

**2.8 ELISA assays**

Norepinephrine (NE) and TNF-α concentration in perfusion heart tissues, perfusate and cell supernatants were determined using the NE research enzyme-linked immunosorbent assay (ELISA) kit (ALPCO#17-NORHU-E01-RES) and the TNF-α Quantikine ELISA kit (R&D

System#RTA00), respectively.

**2.9 Western blotting assay**

Mouse and rat heart homogenates as well as cells were harvested on ice in RIPA lysis buffer (Bioteke#PP1901) containing 1mM phenylmethylsulfonyl fluoride, and then centrifuged at 14,000×g at 4 °C for 15 min. Equal amounts of protein were separated by running 4%–15% SDS-polyacrylamide gel electrophoresis and transferred to PVDF membranes (Millipore#0.45μM). Following blocking with 5% nonfat dry milk for 1 h, the membranes were incubated with the appropriate primary antibodies (described in supplemental material) overnight at 4 °C, followed by incubation with a horseradish peroxidase-conjugated IgG secondary antibody (Dingguo). The immunoreactive bands were visualized with an enhanced chemiluminescence reagent (Millipore, Immobilon$^{TM}$#WBKLS0010), and their intensities were determined by densitometry.

**Antibodies for Western blotting assay**

TH (Abcam#ab112), TLR4 (Abcam#ab22048), DBH (Sigma Aldrich#SAB2701977), P-p65 (CST#3033), P65 (CST#4764), extracellular signalregulated kinase (ERK) 1/2 (CST#4695), P-ERK1/2 (CST#4370), c-jun NH2-terminal kinase (JNK)1/2 (CST#9252), P-JNK1/2 (CST#4668S), p38MAPK (CST#9212S), P-p38 (CST#4511), IκBα (CST#4812S), P-IκBα (CST#9246), glyceraldehyde-3-phosphate dehydrogenase (GAPDH) (CST#2118), c-Fos (CST#2250), c-Jun (CST#9165), CaMKII (CST#3362), P-CaMKII$^{Thr286}$ (CST#12716T), P-CREB$^{Ser133}$ (Affinity#AF3189), CREB (Affinity#AF6188).

**2.10 Quantitative RT-PCR assay**

The mRNA expression was analyzed using standard qRT-PCR protocol. In brief, total RNA was extracted using RNAiso plus reagent (TAKARA#9019) and reverse transcribed using a PrimeScript™ RT Reagent Kit with gDNA Eraser (Perfect Real Time) (TAKARA#RR047A). Real-time PCR were performed with the SYBR Premix Ex Taq II (TAKARA#RR820A) in a LightCycler480 real-time PCR system (Roche#LC480). The expression of each gene mRNA was normalized to that of GAPDH mRNA and results were shown as the fold change to controls. Primers for RT-qPCR were synthesized by Invitrogen™-Thermo Fisher Scientific, Inc (USA). The primer sequences are shown in *Table S3*.

**2.11 Reporter analysis**

293/hTLR4-HA cells seeded on 24-well plates were transiently transfected with 50 ng of the luciferase reporter plasmid together with a total of 300 ng of various expression plasmids or empty control plasmids. As an internal control, pRL-TK was transfected simultaneously. Dual luciferase activity in the total cell lysates was quantified 24-36 h after transfection. Quantification of dual luciferase activity were performed using a dual luciferase assay kit (GALEN#GN201) with standard protocol purchased from YuanPingHao Bio (China).

**2.12 Plasmids and molecular cloning**

pGL3-Basic Luciferase Reporter vector (Promega), pcDNA3.1 plasmid, the plasmids encoding human TRIF and MyD88, RLTK-Luci and AP-1-Luci luciferase reporter plasmid were a gift from Fuping You (Peking University). For reporter assays, rat TH promoter fragments were obtained from neonatal rat genomic DNA by PCR, and then cloned into pGL3-Basic Luciferase Reporter vector with the following promoter regions: −1 to −2000 to get pGL3-Rat-TH-Luci luciferase reporter plasmid. The recombinant plasmid was identified by PCR and restriction enzyme (*Figure S3B-D*). Then the construct was verified by sequencing the relevant region (*Table S4*, *Figure S3E* and *F*). The plasmid encoding EGFP was used as transfection efficiency positive control (*Figure S3G*).

**Reagents for plasmids and molecular cloning**

The reagents genomic DNA extraction kit (#9765), DNA polymerase (LA tag#RR02MA), T4 DNA ligase (#2011A) were purchased from TAKARA. Gel Extracion Kit (#D2500) was purchased from OMEGA bio-tek. Competent cell DH5α (#CD201) was purchased from TransGen Biotech (China), and plasmid DNA extraction kit (#12123) was purchased from QIAGEN (Germany). Restriction enzymes for rapid DNA digestion NheI (#FD0973) and HindIII (#FD0504) were purchased from Thermo Scientific™ (USA). Plasmid transfection was processed using Lipofectamine®3000 Reagent (Invitrogen™#L3000001) and Opti-MEM (Gibco#31985-070).

**2.13 RNA interference**

c-Jun siRNA, c-Fos siRNA (Sequences refer to *Table S5*), and scrambled siRNA, si-h-GAPDH and Cy3-control siRNA were purchased from RIBOBIO (China). siRNA transfection was processed using Lipofectamine®3000 Reagent (Invitrogen™#L3000001) and Opti-MEM (Gibco#31985-070) under standard protocol. RNA interference was designed to disrupt the AP-1 binding in rat TH promoter region. Cy3-control siRNA was used to be a positive control for siRNA transfection (*Figure S4A-C*).

**2.14 CCK-8 assays**

After treatment with drugs, cells were performed cell proliferation assay and cytotoxicity assay using Cell Counting Kit-8 (CCK-8) according to the standard protocol. Briefly, 100 μL of cell suspension was dispensed in a 96-well plate which was pre-incubated in a humidified incubator (5% $CO_2$ at 37°C) for 24 hours. Drugs were then added to the wells. The plate was incubated for an appropriate length of time in the incubator. 10 μL of CCK-8 solution were added to each well of the plate. The plate was incubated for 1-4 hours in the incubator before measured the absorbance at 450 nm using a microplate reader.
